# Different signaling interpretations by PKC eta and theta control T cell function and exhaustion

**DOI:** 10.1101/2024.09.26.615103

**Authors:** Thomas H. Mann, Hannah M. Knox, Shixin Ma, Jesse Furgiuele, Anna-Maria Globig, Michael LaPorta, Hokyung K. Chung, Bryan McDonald, Majid Ghassemian, Steven Zhao, Hubert Tseng, Yagmur Farsakoglu, Victoria Tripple, Johnny Koo, Alexandra C. Newton, Susan M. Kaech

## Abstract

Chronic antigen signaling drives CD8^+^ T cell exhaustion (T_EX_) in cancer and chronic infection. However, how the kinase cascades downstream of the T cell receptor drive exhaustion is not understood. We found that continuous agonism of protein kinase C (PKC) causes degradation of PKC theta, but not PKC eta, and induces terminal T_EX_ cells. During chronic infection, PKC theta is necessary to maintain the progenitor exhausted (T_EX-PROG_) cells, and thus the antigen-specific T cell response, while agonism of PKC eta promotes terminal exhaustion (T_EX-TERM_) *in vitro* and *in vivo*. The cascades downstream of these kinases are distinct, with PKC theta promoting activity of canonical PKC targets in the MAPK and CDK families, while eta promotes activity of other targets, including casein kinase I G2 (CK1G2). Expression of an engineered, degradation-resistant PKC theta, or deletion of the gene encoding CK1G2, improves T cell function and tumor control. Our illustration of multiple therapeutic avenues arising from targeting PKC highlights its centrality in T_EX_ differentiation and its clinical potential in cancer immunotherapy.

**Highlights:**

- PKC theta sustains T cell function while PKC eta promotes terminal exhaustion
- PKC theta and eta drive distinct phospho-cascades to oppose each other’s differentiation instructions
- An engineered, degradation-resistant PKC theta improves T cell responses in chronic infection and cancer
- Ablation of kinase CK1G2 downstream of PKC eta improves anti-tumor T cell responses

## INTRODUCTION

A defining feature of the adaptive immune system is the ability to protect the host by mounting a long-lived response that specifically recognizes foreign antigens. T cells respond to antigen via their T cell receptor (TCR), which initiates kinase cascades and second messenger signaling in response to binding a cognate antigen. Together with costimulatory and inflammatory signals, TCR signals promote activation and differentiation of naïve T cells. Integration of these signals instructs T cells to form diverse cell subsets that can specialize in proliferative capacity, cytotoxicity, and the ability to become tissue-resident. Prior studies of T cells have consequently focused on understanding the different protective capabilities of T cell subsets and on how these subsets form.

During a typical acute infection, T cells give rise to two main subsets of cells, memory precursor (T_MP_) and effector (T_EFF_) cells. T_MP_ are plastic, long-lived cells that produce pro-inflammatory cytokines including IL-2, TNF, and IFNγ. T_MP_ cells seed the pool of memory T cells (T_MEM_) that persists after resolution of the infection to protect from future re-infection by the same pathogen. T_MP_ and T_MEM_ can both undergo extensive proliferation while differentiating to form a spectrum of T_EFF_ cells, which are short-lived but very cytotoxic.

In contrast to the highly functional T_MP_ and T_EFF_ cells that form during acute infections, T cells differentiate into a hypofunctional, ‘exhausted’ set of states during chronic antigen settings such as the response to cancer or HIV. Exhausted T cells (T_EX_) are characterized by their expression of the signature transcription factor TOX, high expression of inhibitory receptors (e.g. PD-1, CTLA-4, TIGIT), and their reduced ability to express pro-inflammatory cytokines upon antigen stimulation. T_EX_ cells exist in a differentiation hierarchy that is a mirror image of their more functional T_MP_ and T_EFF_ counterparts^1,2^. The less-differentiated, lymphoid-resident exhaustion progenitor cells (T_EX-PROG_) self-renewing and, critically, are the cells that proliferate to yield a therapeutic response to anti-PD-1 checkpoint blockade. When T_EX-PROG_ divide, they can then differentiate into either a effector-like cell (T_EX-EFF_) with moderate protective function or a highly dysfunctional, terminally exhausted cell (T_EX-TERM_). Because of the importance of generating and maintaining T_EX-PROG_ and T_EX-EFF_, and avoiding the T_EX-TERM_ state, there has been sustained interest in understanding the balance of T_EX_ subsets. Many signals have been identified that regulate T_EX_ differentiation and function, including cytokines^3^ and hypoxia^4^, among others. But, critically, these environmental factors are not sufficient to induce exhaustion, as antigen signaling is absolutely required to become exhausted: non-specific bystander T cells do not become exhausted even while occupying the same tumor microenvironment as exhausted, antigen-specific cells^5^. Despite ongoing antigen encounter being the prerequisite for T_EX_ formation, how antigen signaling programs T_EX_ differentiation states is poorly understood.

Perhaps the clearest connection of antigen signaling to T_EX_ cell differentiation has been the observation that inhibitory receptor expression is induced by TCR signaling downstream to the transcription factors in the families of nuclear factor of activated T cells (NFAT) and activator protein 1 (AP-1)^6–9^. And studies of inhibitory receptor function have suggested roles for these receptors in regulating the signaling of the TCR and of costimulatory receptors. However, despite the success of immune checkpoint blockade and the central role of antigen signaling in driving T_EX_ formation, few studies have examined the layers of post-translational signaling separating TCR signals from activation of NFAT or other transcription factors (TFs).

T_EX_ cells, and T cell subsets broadly, have largely been defined based on their transcriptional signatures. Indeed, T_MEM_, T_EFF_, and T_EX_ all have distinct transcriptional profiles, and studies of the signature TFs regulating these states have been elucidated much of how they function. Unique epigenetic profiles regulate these transcriptional states^10,11^, including driving a commitment to the T_EX_ lineage that cannot be reversed by immune checkpoint blockade^12,13^. These studies have yielded a rich view of differentiation state-specific transcription machinery, but meanwhile, studies of TCR signaling have largely been conducted in homogenous populations of cell lines or naïve T cells that are undergoing priming. This contrast lays bare several major questions. Do T_EX_ cell subsets signal differently downstream of the TCR? How does subset-dependent signaling regulate T_EX_ differentiation and function? And what are the therapeutic implications of any differences?

Here, we found that CD8^+^ T_EX_ cell subsets engage distinct signaling pathways downstream of the TCR by employing two different members of the protein kinase C (PKC) family, PKC theta and PKC eta. These two kinases had opposing effects on T_EX_ differentiation, as PKC theta supported T_EX-PROG_ maintenance and T cell effector functions while PKC eta promoted T_EX-TERM_ formation. In viral and tumor models of T cell exhaustion, we found that PKC theta was selectively targeted for proteolytic degradation upon its stimulation while PKC eta was maintained. We found that these kinases drive distinct phosphoproteomes, which in turn lead to different cell fates. Our study demonstrates that T cell fates are intertwined with the antigen receptor signaling capacities of that cell state and further reveals that the differential abundance and functions of the PKCs are key to this relationship. This finding illustrates how kinases can act as lineage determining factors in T cells and highlights how they can be manipulated in the T cell response to cancer and viruses.

## RESULTS

### PKC theta is required for T_EX_-_PROG_ while PKC eta promotes T_EX-TERM_ differentiation

CD8^+^ T cells have been defined as exhausted based on their reduced ability to produce proinflammatory cytokines or to kill target cells presenting their cognate antigen, but little has been done to investigate how signaling downstream of the TCR is intertwined with T_EX_ cell fates. To test if there are differences in kinase signaling between T_EX_ subsets, we restimulated T_EX_ cells from LCMV Clone 13 (Cl 13) infected mice 30 days post-infection (d.p.i.) *ex vivo* with phorbol myristate acetate (PMA), which agonizes protein kinase C proteins by mimicking the endogenous ligand, diacylglycerol, and analyzed phosphorylation of proteins by phospho-flow cytometry. While some targets downstream of PKC, such as ERK (p44/42) and NF-kB p65, were restimulated equally well between SLAMF6^+^ TIM3^-^T_EX-PROG_ and SLAMF6^-^ TIM3^+^ CD101^+^ T_EX-TERM_ cells, p38, JNK, and c-Fos had lower amounts of phosphorylation in T_EX-TERM_ cells (**Figure 1A**). These discrepancies following PMA restimulation, indicated that PKC signaling differs between the T_EX-PROG_ and T_EX-TERM_ subsets. Indeed, we found that T_EX_ subsets expressed different amounts of two PKC proteins, PKC theta and eta, the dominant PKCs expressed in CD8^+^ T cells. PKC theta expression was highest in T_EX-PROG_, maintained in T_EX-EFF_, but then decreased in T_EX-TERM_ cells, while PKC eta showed an opposite trend, with protein levels highest in T_EX-TERM_ cells; a similar trend for PKC theta levels was observed in T_EX_ subsets isolated from MC38 tumors (**Figure 1B** and **Figure S1A**). The expression patterns of PKC theta and eta were also observed in published mRNA data^14^, correlating with alterations in the expression of genes in the AP-1 family, such as c-Jun (**Figure S1B-C**). Additionally, the mRNA expression of other PKCs, such as alpha, beta, and delta, was limited relative to PKC theta and eta, leading us to focus our study on these two proteins (**Figure S1B**).

**Figure 1.**
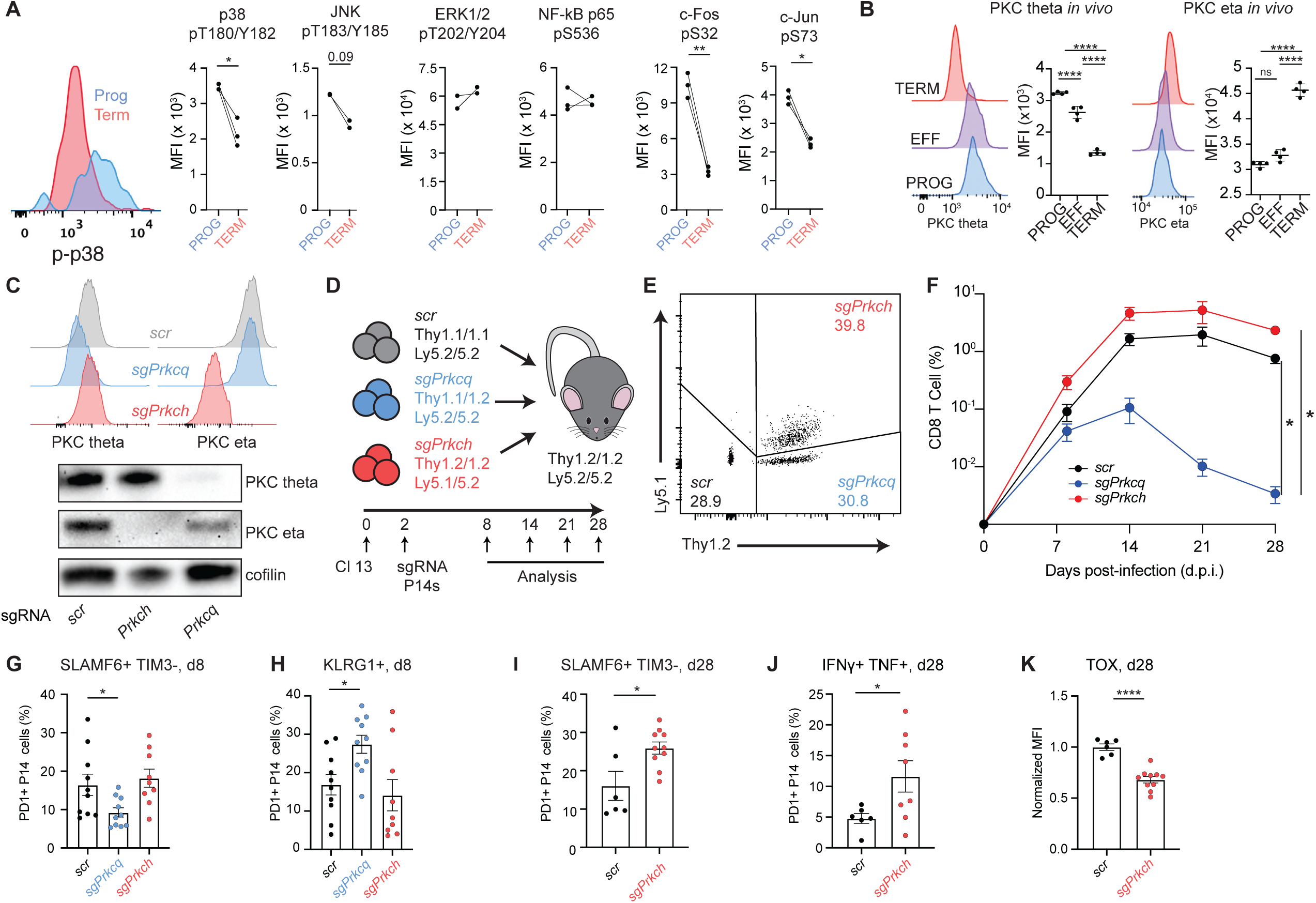
Distinct kinase activities of PKC theta and eta reflect opposing functions in CD8^+^ T cell differentiation. (A) MFIs from phospho-flow cytometry of the indicated markers are shown. T_EX-PROG_ or T_EX-TERM_ cells from LCMV Cl 13 infected mice 30 d.p.i. were stimulated with PMA *ex vivo* for 30 mins (all but c-Fos and c-Jun) or 4 hours (c-Fos and c-Jun). (B) Expression of PKC theta and eta by subset from LCMV Cl 13, 30 d. p.i. (C) Validation of PKC theta (*Prkcq*) and PKC eta (*Prkch*) knockouts by flow cytometry and Western blot. (D) Mice were infected with LCMV clone 13, and congenic P14 T cells were negative selected and activated *in vitro* via platebound anti-CD3/28. After 24 hours, cells were subjected to Cas9-sgRNA electroporation to delete genes of interest, and after a further 24 hours of recovery, electroporated cells were mixed and co-transferred into infected mice (E-F) The frequencies of each congenic population of electroporated cells at the time of co-transfer (E), and the frequencies of cells of each genotype relative to the total CD8 T cell population at the indicated timepoints post-infection (F) (G-H) The frequencies of indicated markers among transferred PD1+ CD8 T cells, 8 d.p.i (I-K) The frequencies of indicated markers (I-J) and TOX MFI (K) in transferred PD1+ cells, 28-30 d.p.i Scatter plots show mean +/- s.e.m. Statistical significance was determined using Student’s t-test (A, I-K) or one way ANOVA (panels B-H), with significance shown as *P < 0.05, **P < 0.01, ***P < 0.001, ****P < 0.0001.

To test the roles of PKC theta and eta in CD8^+^ T cell differentiation, we established a system for deleting *Prkcq* or *Prkch* (coding for PKC theta and eta, respectively) in virus-specific T cells using CRISPR/Cas9^15^. We electroporated P14 CD8^+^ T cells, specific for the gp33 peptide from LCMV, with Cas9 ribonucleoprotein (RNP) complexes containing guide RNAs against the PKC genes, or an untargeted scramble, “scr,” control and validated the deletions by flow cytometry and Western blot (**Figure 1C**). Because of PKC theta’s well-established role in activation of naïve T cells, these genes were deleted in P14 cells 24 hours after *in vitro* activation. Another 24 hours later (48 hours since initial activation), we then mixed *scr*, *sgPrkcq*, or *sgPrkch* P14 cells in approximately equal numbers and adoptively transferred this mixture into mice that had been infected 48 hours prior with LCMV Cl 13 (**Figure 1D-E**). Notably, P14 cells lacking *Prkcq* precipitously declined in numbers over the course of infection from 14 d.p.i. onward, while the kinetics of cells lacking *Prkch* resembled those of the WT control *scr* cells (**Figure 1F, Figure S1D-E**).

To understand whether differentiation changes caused by deletion of *Prkcq* could explain the antigen-specific cell population collapse, we examined markers of differentiation at 8 d.p.i., when the *sgPrkcq* population remained viable. Indeed, at this timepoint, we found that *sgPrkcq* cells were impaired in their ability to form the long-lived T_EX-PROG_ cells that maintain the Ag-specific response (**Figure 1G**). Instead, they more frequently expressed KLRG1, a marker of differentiation into terminal effector (TE) cells (**Figure 1H**), a population that readily undergoes apoptosis^16^. The effects of deleting *Prkcq* in CD8^+^ T cells resembled those observed for deletion of *Tox*^17,18^ or *Tcf7*^19^ because more KLRG1^+^ terminal effector-like cells formed at the expense of T_EX-PROG_ cells and the pool of virus-specific CD8^+^ T cells was not maintained over time. Notably though, this phenotype occurred by deletion of a kinase involved in TCR and costimulatory signaling, rather than a lineage-specifying TF.

In contrast to *sgPrkcq* cells, deletion of *Prkch* in P14 CD8^+^ T cells led to an increase in the frequency of T_EX-PROG_ cells at 28 d.p.i. (**Figure 1I**). Additionally, these cells exhibited increased production of IFNγ and TNF upon restimulation and expressed lower amounts of the exhaustion TF TOX (**Figure 1J-K**). Altogether, these results demonstrate that PKC theta and eta have opposing functions in CD8^+^ T cell differentiation, namely that PKC theta promotes T_EX-PROG_ while PKC eta promotes T_EX-TERM_ cell formation.

### Chronic agonism of PKC *in vitro* is sufficient to drive a terminal exhaustion signature in CD8^+^ T cells

To better dissect the regulation of PKC signaling in activated CD8^+^ T cells we turned to *in vitro* exhaustion strategies used by other groups^20,21^ and continuously co-cultured P14 T cells with B16-gp33 tumor cells for a week to progressively induce T cell dysfunction with or without the PKC agonist PMA to see if this affected *in vitro* T cell exhaustion (**Figure 2A**). Strikingly, inclusion of PMA in these *in vitro* exhaustion cultures potently exacerbated features of terminal T cell exhaustion, including loss of expression of TCF1, increased expression of TIM3 and TOX, and reduced production of IFNγ and TNF production (**Figure 2B-C**). Similar results were obtained using another PKC agonist, bryostatin (**Figure S2A**). PMA stimulation of activated CD8^+^ T cells *in vitro* also induced a T_EX-TERM_ gene expression signature including *Tox*, *Havcr2*, *Pdcd1*, and *Gzmb* (**Figure 2D**). These data indicated that chronic agonism of PKCs *in vitro* was sufficient to drive cardinal features of terminal exhaustion.

**Figure 2.**
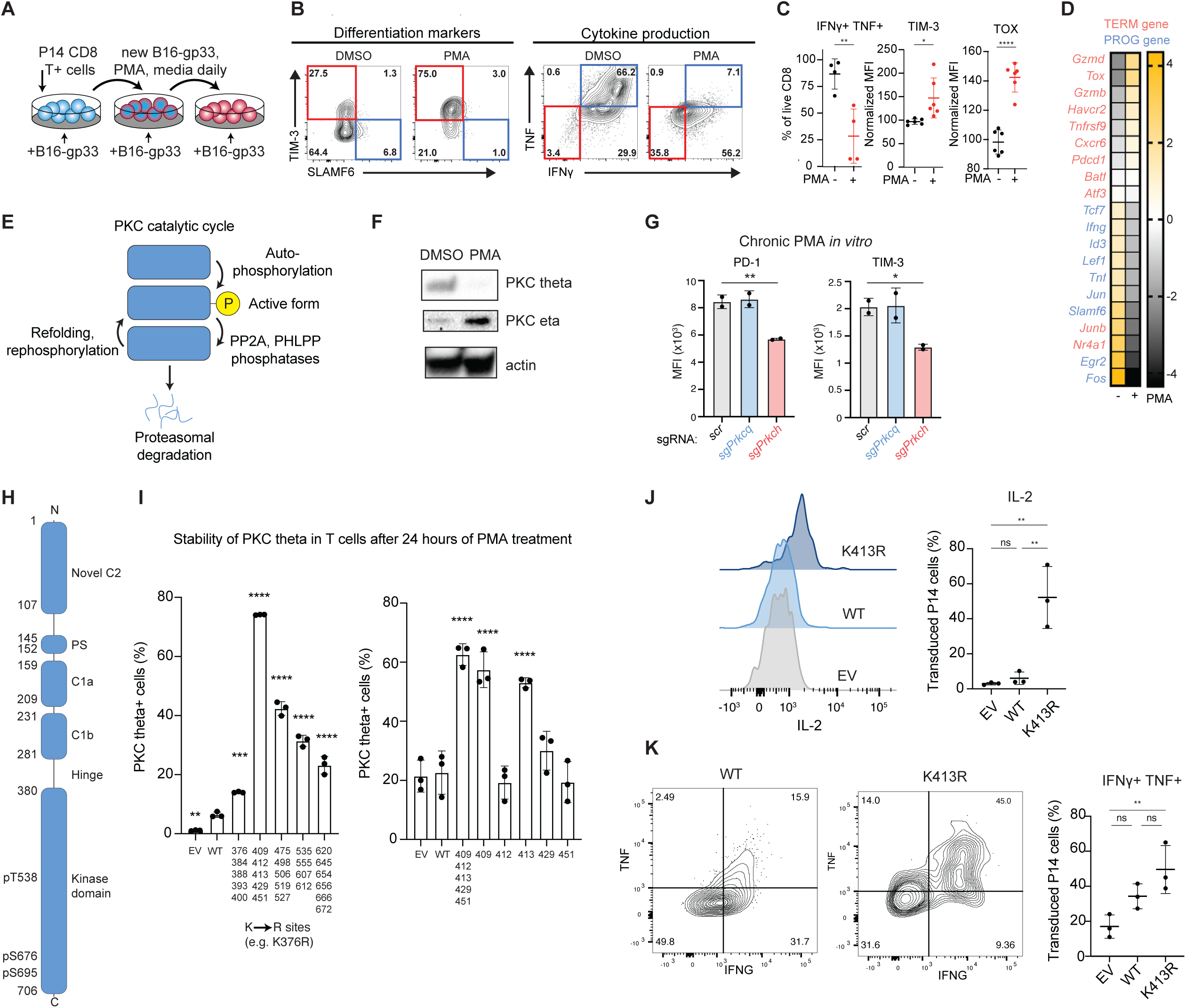
Chronic agonism of PKC drives terminal exhaustion in CD8+ T cells by promoting PKC theta degradation, which can be rescued by mutation of K413. (A) Schematic of *in vitro* exhaustion experiments. P14 T cells were activated *in vitro*, expanded, and co-cultured with B16-gp33 cells from days 3-7 post-activation. (B) Representative flow plots showing the effect of chronic PMA treatment on T cells *in vitro*. (C) Cumulative bar graphs of PMA effects on *in vitro* exhausted cells. (D) nCounter data highlighting PMA-driven gene expression changes. T_EX-TERM_ associated genes are in red, while T_EX-PROG_ associated genes are in blue. Data are shown as log2 fold change. (E) Cartoon schematic of the regulation of PKC kinase activity. (F) Western blots showing PKC protein levels in cultured primary CD8^+^ T cells after 4 days of PMA treatment. (G) Effects of KO of *Prkcq* or *Prkch* on responsiveness to PMA during *in vitro* exhaustion. (H) Domain architecture of PKC theta. Numbers represent the amino acids defining each domain and the three phosphorylation sites key for PKC kinase activity. (I) A two-round K-to-R mutagenesis screen to identify lysine residues required for PKC activity-induced degradation after 24 hours of PMA treatment. Lysine residues were first mutated in blocks of four to six at a time (left), and then individual lysine residues were mutated within the 409-451 subset (right) (J-K) P14 T cells were transduced with EV or PKC OE vectors, subjected to *in vitro* exhaustion, and assessed for their capacity to produce cytokines. Scatter plots show mean +/- s.e.m. Statistical significance was determined using Student’s t-test (panel C) or one way ANOVA (panels G, I-K), with significance shown as *P < 0.05, **P < 0.01, ***P < 0.001, ****P < 0.0001.

### Chronic agonism of PKCs drives degradation of PKC theta, while eta is maintained

The observation that PKC theta and eta have different genetic effects on T cell differentiation *in vivo* provides an intriguing contrast with the finding that continuous agonism of both PKCs by PMA treatment was sufficient to drive features of T_EX-TERM_ cells *in vitro* (**Figure 2A-D**). We hypothesized that a resolution of this paradox would be if chronic stimulation had different effects on PKC theta and eta activity or stability.

PKCs become activated and deactivated in a multi-step cycle that includes the potential for proteasomal degradation (**Figure 2E**)^22^. PKCs naturally exist in an autoinhibited state and must be phosphorylated to become active towards their substrates. Upon dephosphorylation, by the PP2A or PHLPP phosphatases, a PKC molecule can then either be protected by Hsp70 to refold into the primed but inactive state, or alternatively be targeted by E3 ligases to drive proteasomal degradation. We hypothesized that PKC theta or eta could differ in their regulation by one of these feedback mechanisms to induce T_TERM_ phenotypes when both are stimulated.

Because of the central role of proteasomal degradation in PKC regulation, we first tested if chronic PMA stimulation affected PKC theta and eta protein levels differently. Prior studies of EL-4 cells had indeed shown that PKC eta was selectively maintained and upregulated during chronic PKC stimulation^23^. Notably, in primary CD8^+^ T cells, Western blotting showed that PKC theta was completely degraded after several days while PKC eta was maintained (**Figure 2F**). Subsequent kinetics showed that PKC theta degradation was complete within 24 hours, and that Hsp70 inhibition accelerated this process (**Figure S2B-E**). The selective maintenance of PKC eta but not theta after chronic PMA stimulation is significant in that it mimics the PKC expression pattern as T cells differentiate from T_EX-PROG_ to T_EX-TERM_ *in vivo* (**Figure 1B**).

Given that PKC eta, but not theta, was present during chronic PMA stimulation during *in vitro* exhaustion assays, we tested whether PKC eta was necessary for the PMA-induced effects on exhaustion. We transduced Cas9-expressing P14 T cells with sgRNA-encoding retroviral vectors against each PKC, and we found that PKC eta was required for maximal induction of PD- 1 or TIM3 during PMA treatment (**Figure 2G**). These findings show that agonizing PKC eta *in vitro* can induce key features of T cell exhaustion, and that PKC eta was necessary for T cells to acquire these features because it is selectively maintained relative to PKC theta.

### Engineering a degradation-resistant PKC theta by scanning mutagenesis of lysine residues

Because PKC theta was rapidly degraded during periods of intense stimulation and this correlated with acquisition of T_EX-TERM_ cell phenotypes and impaired cytokine production, we hypothesized that preventing PKC theta degradation may result in superior CD8^+^ T cell function by sustaining effector functions or T_EX-PROG_ states. To test this idea, we sought to create a degradation-resistant form of PKC theta.

E3 ligases target proteins for degradation by fusing ubiquitin to lysine residues^24^. The E3 ligase PELI1 was previously shown to ubiquitylate PKC theta^25^, but the lysine(s) it targets was unknown. Deletion of *Peli1* had little effect on amounts of PKC theta in T_EX-PROG_ cells but led to a significant increase in PKC theta protein levels in T_EX-EFF_ cells and trended upwards in T_EX-TERM_ cells too, despite their lower transcription of *Prkcq* (**Figure S2F**).

We found that the kinase domain (KD), at the C-terminus of PKC (**Figure 2H**), could be degraded during chronic PMA treatment (**Figure S2G**). To identify the lysine residue(s) responsible for degradation, we designed PKC variants in which we systematically mutated every lysine residue in the KD, first scanning the protein as five bins containing 4 to 6 lysine residues each (**Figure 2I**). Within each bin, the lysines were mutated to arginine to preserve a positive charge while blocking ubiquitination. We next transduced these PKC theta variants along with WT PKC theta and empty vector (EV) controls into activated CD8^+^ T cells, stimulated the cells with PMA for 24 hours, and then measured the abundance of each PKC. As expected, PMA treatment was sufficient to induce degradation of overexpressed WT PKC theta. However, one variant (Bin 2, encoding mutations K409R, K412R, K413R, K429R, and K451R) exhibited PKC theta abundance far above the other constructs, though several other variants showed partial effects on PKC theta levels during prolonged PMA stimulation. To narrow down if there were specific dominant lysine residues regulating PKC theta degradation in Bin 2, we proceeded to mutate each residue individually to arginine. Two mutations, K409R and K413R, occurring in a region of near-perfect amino acid conservation between mouse and human (**Figure S2H**), displayed similar degradation resistance as the entire Bin 2 mutant variant. However, K409R has previously been identified as a catalytically dead version of PKC theta: K409 contacts ATP in the binding pocket, and this mutation interferes with ATP binding^26^. Meanwhile, K413 points out into solution in the structure of PKC theta, suggesting it as a ubiquitination target (**Figure S2H**)^27^.

To test whether the degradation resistant PKC theta K413R improved T cell function, P14 cells were transduced with PKC theta K413R and the cells were cultured under strenuous *in vitro* exhaustion conditions, including chronic PMA treatment for 4 days (**Figure 2A-D**). Indeed, PKC theta K413R transductants displayed dramatically increased IL-2, IFNγ, and TNF production. (**Figure 2J-K**). Altogether, these results show that expression of a degradation-resistant variant of PKC theta is sufficient to maintain high levels of T cell effector function even during periods of intense, prolonged stimulation.

### Overexpression of PKC theta K413R drives improved T cell function *in vivo*

We hypothesized that overexpression of PKC theta, particularly the degradation-resistant PKC theta K413R, would improve anti-viral and anti-tumor CD8^+^ T cell responses *in vivo*. To test this idea, first, P14 cells expressing PKC theta WT, K413R, or an EV control were adoptively transferred into mice infected with LCMV Cl 13 and analyzed 28-30 d.p.i. (**Figure 3A**). While the overall amounts of PKC theta were not significantly changed across the whole population of GFP^+^ transductants, they were significantly increased in T_EX-TERM_ cells OE PKC theta, particularly for PKC theta K413R (**Figure 3B-C**). This pattern reflects that PKC theta is targeted for degradation as T_EX-TERM_ cells develop during chronic infection. OE of PKC theta WT trended to increase the total number of GFP^+^ transductants recovered from the spleen, but OE of degradation-resistant PKC theta K413R led to a striking tenfold increase in numbers of virus-specific CD8^+^ T cells (**Figure 3D**). Moreover, the two PKC theta OE vectors, WT and K413R, differed in their capacities to boost individual cell subsets. Both WT and K413R OE increased the number of SLAMF6^+^ TIM3^-^ T_EX-PROG_ cells to similar extents relative to EV. But K413R was unique in dramatically increasing the number of SLAMF6^-^ TIM3^+^ cells, which includes both T_EX-EFF_ and T_EX-TERM_ cells (**Figure 3E-F**). However, other measures of T cell state and effector function were little changed (**Figure 3G-I**), indicating that *in vivo*, during a chronic infection the primary effect of OE PKC theta, particularly K413R, is to yield a much larger antigen-specific CD8^+^ T cell population.

**Figure 3.**
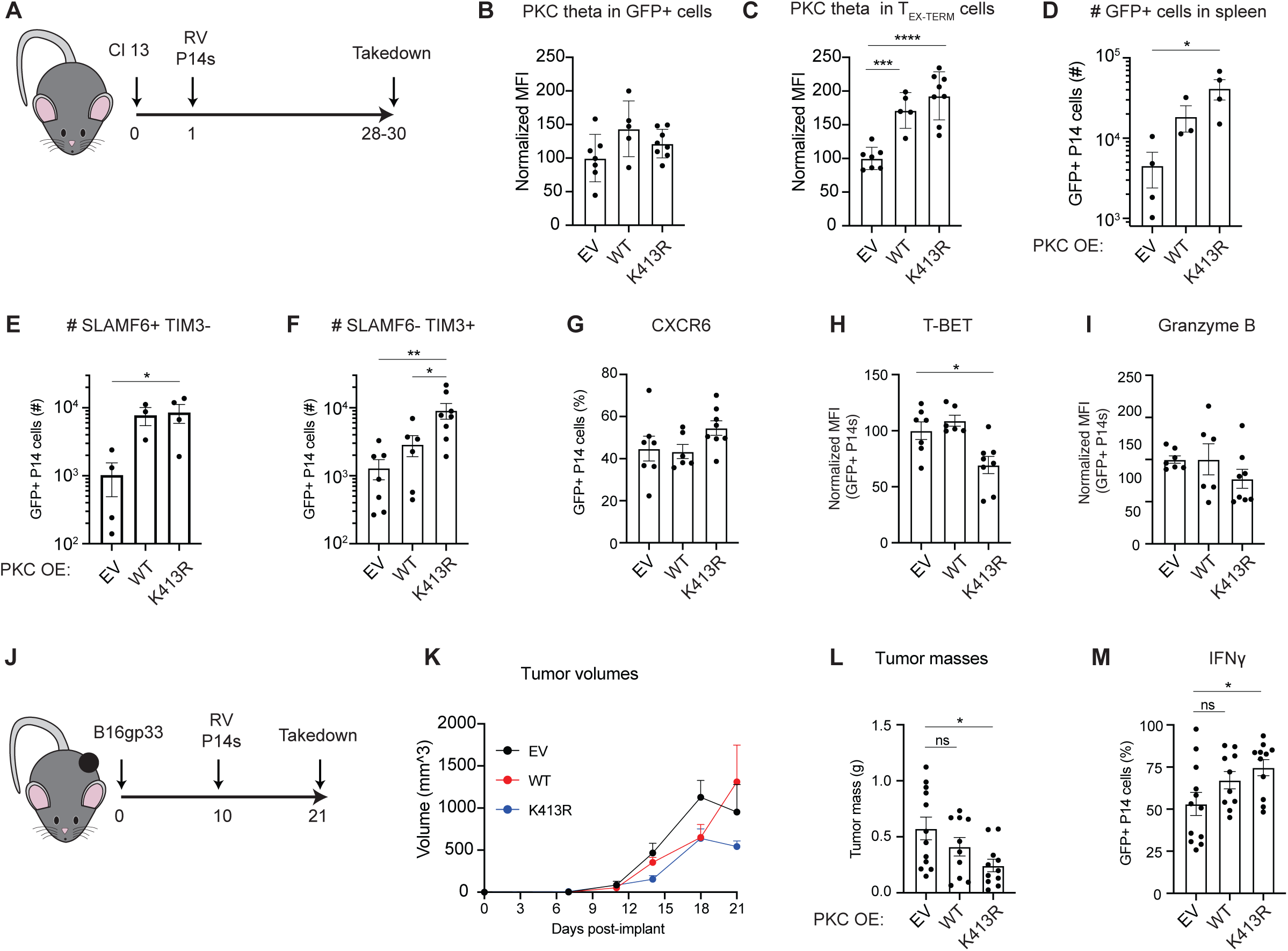
Overexpression of degradation-resistant PKC theta promotes anti-tumor function *in vivo*. (A) Timeline of adoptive transfer experiments of P14 T cells into mice infected with LCMV Cl 13. Overexpression of PKC theta variants or an EV control were introduced using a retroviral (RV) vector. (B-C) MFI of PKC theta among all transferred GFP+ (B) and GFP+ T_TERM_ cells (C). (D-F) Numbers of total GFP+ cells of the indicated genotypes, in total (D), or in SLAMF6^+^ TIM3^-^ (E) or SLAMF6^-^ TIM3^+^ (F) subsets. (G) Frequency GFP+ P14 T cells expressing CXCR6 in LCMV Cl 13. (H-I) Normalized MFI of the indicated markers in LCMV Cl 13. (J) Timeline of adoptive transfer experiments in B16gp33 tumor-bearing mice. (K-L) Tumor volumes (K) and masses at endpoint (L) of tumors by P14 genotype. (M) IFNγ producing functionality of P14 T cells infiltrating B16gp33 tumors by genotype. Scatter plots show mean +/- s.e.m. Statistical significance was determined using one way ANOVA where applicable, with significance shown as *P < 0.05, **P < 0.01, ***P < 0.001, ****P < 0.0001.

Second, to further probe the potential therapeutic efficacy of OE of PKC theta WT and K413R, P14 cells overexpressing these constructs or EV controls were transferred into B16gp33 tumor-bearing mice at d10 post-tumor implant (**Figure 3J**). This showed that OE of PKC theta K413R provided effective suppression of tumor growth, while cells transduced with WT PKC theta or EV failed to control tumors (**Figure 3K-L**). Additionally, P14 T cells expressing PKC theta K413R produced more IFNγ (**Figure 3M**). Together, these data indicate that OE of PKC theta, particularly a degradation-resistant form, improves CD8^+^ T cell responses in chronic viral infection and in tumors. The sequence conservation around K413 between mice and humans (**Figure S2H**) suggests a high translational relevance. These findings underscore that the loss of expression and proteolytic degradation of PKC theta are key steps in the progression of T cell exhaustion and that artificially maintaining PKC theta is sufficient to boost T cell effector function.

### Phospho-proteomics reveal different kinase targets for PKC theta and eta

The finding that deletions of the genes encoding PKC theta and eta have opposing effects on T_EX_ cell differentiation — and that PKC theta is more abundant in T_EX-PROG_ cells whereas PKC eta is more abundant in T_EX-TERM_ cells — suggests that these kinases execute separate downstream signaling programs to regulate cell fate. To elucidate the changes in signal transduction that dictate T cell exhaustion and connect these opposing functions in T_EX_ cells, we performed global phosphoproteomics on primary CD8^+^ T cells lacking PKC theta or eta (**Figure 4A**). CD8^+^ T cells were electroporated with sgRNAs against each PKC or a non-targeting guide 24 hours after their platebound activation with anti-CD3/28. After four days of expansion, the effects of anti-CD3 stimulation were tested in each background by restimulating half of the cells of each genotype with anti-CD3 for 30 minutes, and pellets of the stimulated or unstimulated cells were collected for phosphoproteomics.

**Figure 4.**
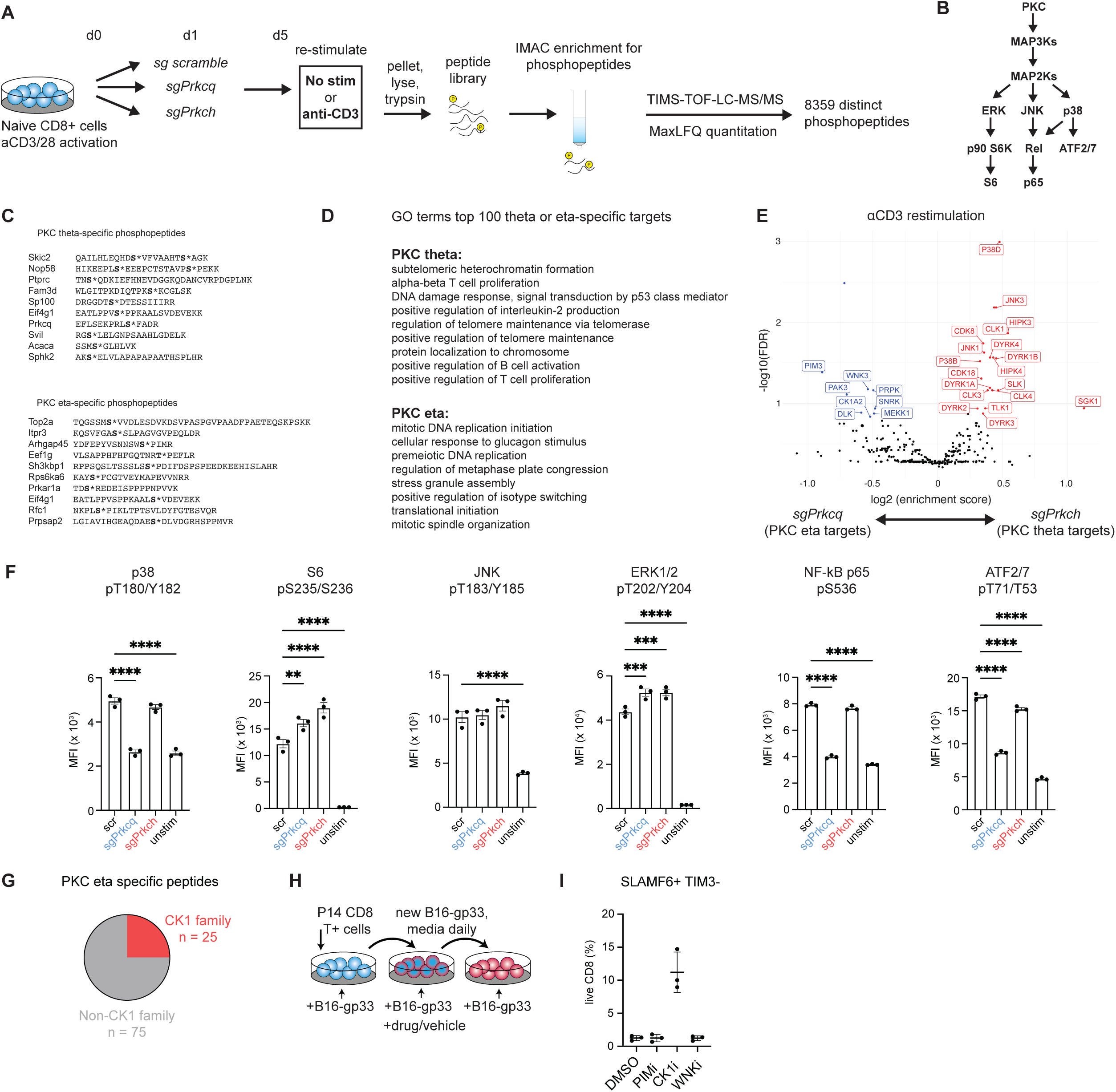
PKC theta preferentially activates the MAPK and CDK pathways, whereas PKC eta activates distinct phosphorylation cascades, including CK1. (A) Schematic of preparation of primary T cells for global phosphoproteomics. (B) Cartoon highlighting and simplifying major known pathways downstream of PKC. (C) Select, top-ranking PKC theta-specific or PKC eta-specific phosphopeptides identified from the anti-CD3 stimulation condition. Phosphorylated residues are highlighted in bold and followed with an asterisk (*). The gene name of the protein from which the peptide originates is shown to the left of each peptide. (D) GO terms associated with the top 100 PKC theta- vs eta-specific downstream phosphoproteins. (E) Volcano plot of Kinase Library analysis of kinases active downstream of PKC theta or eta after anti-CD3 stimulation. Re-stimulated *sgPrkch* or *sgPrkcq* samples were normalized to unstimulated controls of the matching genetic background, and the normalized phosphoproteomes were subsequently compared to one another. The kinases active in each condition (plotted) are inferred by identifying enrichment of ideal peptide motifs for each kinase in the entire proteomics dataset. (F) Phospho-flow cytometry of the indicated markers within *in vitro* activated CD8 T cells, stimulated for 30 minutes with PMA, in the indicated genetic backgrounds. (G) Fraction of top 100 PKC eta-specific peptides that score highly as CK1 family targets according to Kinase Library peptide motifs. (H) Schematic of *in vitro* exhaustion experiments used to screen inhibitors of kinases downstream of PKC theta and eta. (I) Screening effects of inhibitors of kinase families putatively downstream of the PKCs for markers of T_EX-PROG_ differentiation. Scatter plots show mean +/- s.e.m. Statistical significance was determined using one way ANOVA where applicable, with significance shown as *P < 0.05, **P < 0.01, ***P < 0.001, ****P < 0.0001.

Kinases frequently function in multi-step cascades which enable signal amplification at each level, crosstalk between pathways, and for individual components to have distinct subcellular or temporal regulation. PKC enzymes generally regulate the mitogen activated kinase (MAPK) and cyclin-dependent kinase (CDK) pathways to regulate cell growth, proliferation, and certain effector functions. In T cells, PKC theta is known to be an important driver of cytokine production through its activation of AP-1 and nuclear factor kappa-light-chain-enhancer of activated B cells NF-kB (**Figure 4B**). However, because these pathways do not necessarily represent the entire set of targets regulated by PKC theta and eta, phosphoproteomic analysis of can provides a more comprehensive view of their activity.

To gain a broader view of the distinct phosphoproteomes regulated by PKC theta or eta, we compared the abundances of individual phosphopeptides generated following anti-CD3 stimulation in cells lacking either PKC theta or eta. Following label-free quantitation^28^, peak areas of individual peptides were normalized in each anti-CD3 stimulated condition to their abundance in unstimulated cells of the same genetic background (i.e. stimulated *sgPrkcq* cells were normalized to unstimulated *sgPrkcq* cells). The log2 fold changes of individual peptides were then calculated between *sgPrkcq* and *sgPrkch* conditions and ranked according to the largest fold changes (**Figure 4C**). Among these peptides, a phosphopeptide containing PKC theta phosphorylated at S676, an autophosphorylation indicative of PKC theta activity, ranked as one of the most enriched peptides in *sgPrkch* cells compared to *sgPrkcq* cells, validating the overall quantitation strategy. Gene ontology (GO term) analysis of the proteins containing the top 100 PKC theta or eta-specific peptides reflected differences in the biology of these enzymes (**Figure 4D**). While PKC theta was clearly associated with multiple T cell-specific functions, PKC eta was associated with more general cellular processes, including chromosome regulation and stress response pathways.

We hypothesized that PKC theta and eta may enact distinct effects on T_EX_ cell fate by regulating different branches of classic PKC target pathways, or by regulating different pathways entirely. To globally assess the kinases active downstream of PKC theta and eta, the Kinase Library^29,30^ was used to analyze differential phosphoproteomics data. Using established peptide motifs preferred by hundreds of mammalian kinases, the Kinase Library identifies which motifs are enriched in phosphoproteomics datasets and thus predicts which kinases are active in a particular condition, similar to TF motif analysis by HOMER^31^.

When we directly compared the PKC theta-dependent phosphoproteome to that of PKC eta, we found that PKC theta preferentially activates multiple canonical PKC target proteins, including p38, Jun N-terminal kinase (JNK), and several CDKs (**Figure 4E**). Meanwhile, kinase families dependent upon PKC eta for activation included Pim, p21 activated kinase (PAK), with no lysine/K (WNK), and casein kinase 1 (CK1). Aside from a CTLA-4-dependent interaction between PKC eta and PAK2 in Treg cells^32^, PKC eta has not previously been associated with these kinases. These findings are somewhat surprising, as similarities between PKC theta and eta domain architectures and target peptide motifs would initially suggest overlapping functions. However, the propensity of N-terminal sensory domains in PKCs to dictate their subcellular localization and protein interaction partners^33,34^ provides a potential mechanism for the different pathways putatively activated by PKC theta and eta.

To validate the PKC theta and eta-target differences suggested by phosphoproteomics, primary T cells were electroporated with guide RNAs against *Prkcq*, *Prkch*, or a scr control. These cells were expanded for several days and restimulated for 30 minutes either with PMA to specifically agonize PKC (**Figure 4F**), and phospho-flow cytometry was used to measure the activation of several proteins downstream of the PKCs. Within the MAPK pathway (**Figure 4B**), PKC theta and eta were equally able to activate the ERK/pS6 portion of the cascade. PKC theta was required for activation of other portions of the pathway, including p38, NF-kB p65, and ATF2/7, while ablation of PKC eta had no effect. Broadly, the phosphoproteomic and phospho-flow cytometric data indicate that PKC theta and eta have distinct signaling fingerprints which likely drive their differential regulation of T_EX_ cell fates.

To validate the PKC eta targets suggested by phosphoproteomics, we began by scoring the top 100 PKC eta-specific phosphopeptides as potential substrates for different kinases. Among these, 25% of them scored highly as candidate CK1 targets (**Figure 4G**). So, we tested if inhibiting these kinases and others putatively downstream of PKC eta would maintain T_EX-PROG_ markers on T cells subjected to our *in vitro* exhaustion assay (**Figure 4H**). The pan-CK1 inhibitor, D4476, indeed maintained cells in a SLAMF6^+^ TIM3^-^ state *in vitro*, suggesting that PKC eta may activate CK1 to repress T_EX-PROG_ cell differentiation (**Figure 4I**). Altogether, these data show that PKC theta and eta activate distinct signaling cascades and that some of these PKC eta targets like CK1 may promote T cell exhaustion. PKC theta preferentially activates canonical PKC substrates in the MAPK and CDK pathways, while PKC eta promotes the activity of other kinase families, including CK1, WNK, PIM, and PAK.

### PKC eta signals through pathways independent of the PKC theta-CARMA1 axis

To better understand how PKC theta and eta may establish distinct downstream signaling modules, we explored their protein interaction partners. PKC proteins typically signal by phosphorylating a scaffold protein, which then becomes competent to recruit substrates for PKC to phosphorylate. While the signaling scaffold for PKC theta is well described^35^ (comprised of CARMA1, BCL10, and MALT1, the CBM complex, **Figure 5A**) and essential for normal T cell activation and cytokine production^36,37^, the scaffold for PKC eta is unknown. To test whether PKC eta also signals through the canonical CBM complex, we activated primary CD8^+^ T cells, deleted *Prkcq* and *Prkch,* and later treated the cells with vehicle or PMA for 30 minutes to stimulate PKC activity. As predicted, PKC theta was required for cleavage of the MALT1 substrate Regnase-1^38^, but deletion of PKC eta had little effect on Regnase-1 cleavage (**Figure 5A-B**), indicating that PKC eta poorly activates the canonical CBM complex. This result indicates that PKC eta mediates distinct downstream activities from PKC theta by interacting with different scaffold complexes, potentially at different subcellular locations

**Figure 5.**
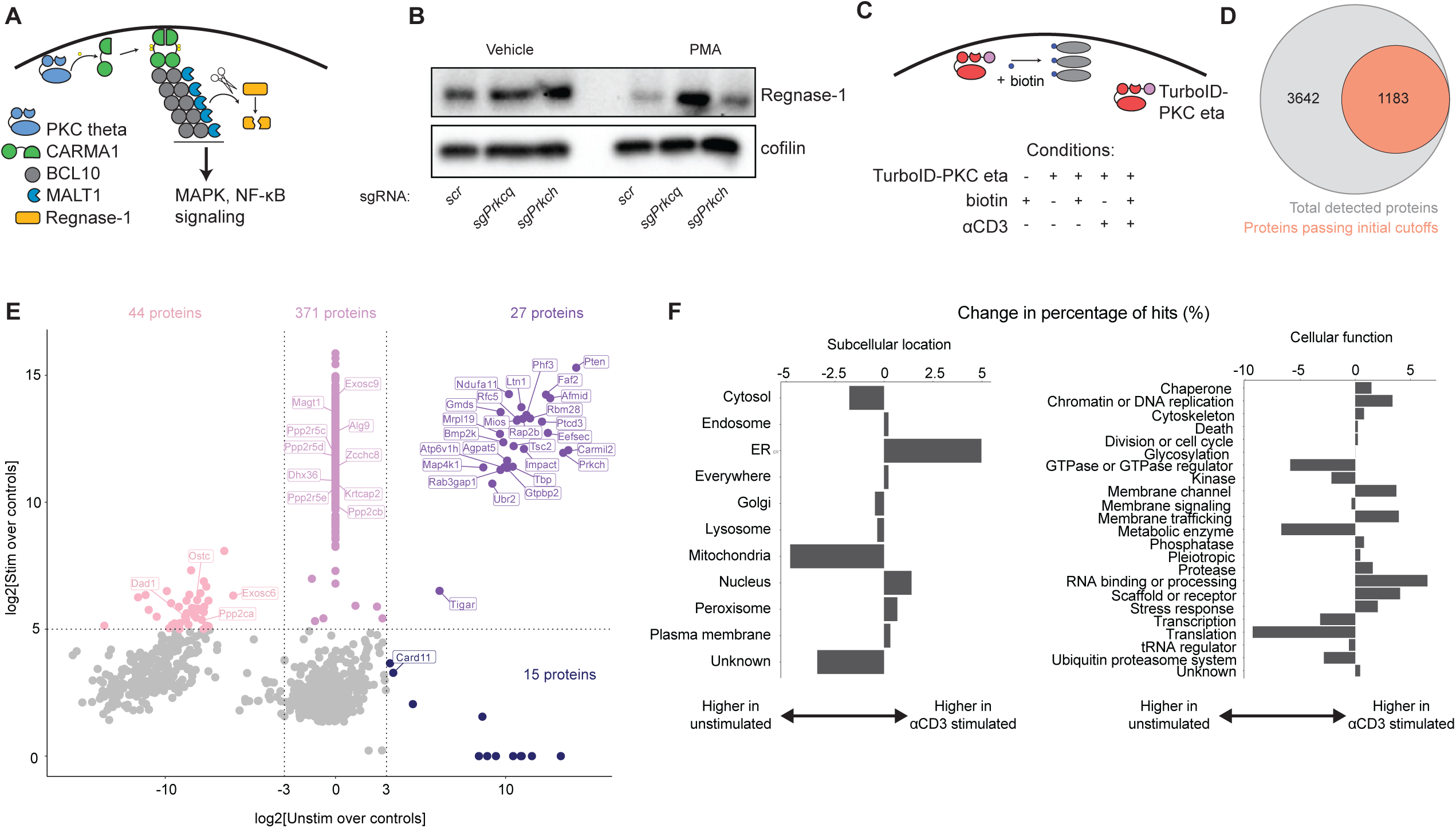
PKC eta interacts with proteins throughout the cell to signal independently of the PKC theta scaffold complex CARMA1-BCL10-MALT1. (A) Cartoon of PKC theta signaling through the CARMA1-BCL10-MALT1 complex. (B) Western blot showing Regnase-1 levels after 30 minutes of PMA or vehicle treatment. T cells were collected after *in vitro* activation and 5 days of expansion. (C) Cartoon schematic of PKC eta proximity labeling strategy and conditions via TurboID. The TurboID-PKC eta fusion protein was transduced into primary T cells, and the expanded transductants were treated and collected 6-7 days post-activation. (D) Venn diagram of raw TurboID statistics. (E) Scatterplot of comparing the log2 fold enrichment of candidate PKC eta interacting proteins in anti-CD3-stimulated vs unstimulated conditions. Dashed lines and colors indicate protein subsets based on enrichment thresholds. Enrichment value was determined by dividing the peak area of candidate proteins by the mean peak area of negative control conditions. Only proteins with p_adj_ < 0.01 are plotted. (F) Changes between the subcellular localizations and cellular functions of top candidate PKC eta interactors between stimulated and unstimulated conditions.

Because the scaffold of PKC eta has not yet been characterized, we performed TurboID^39^ analysis of PKC eta to identify its interaction partners. P14 CD8^+^ T cells were transduced with TurboID-PKC eta or an EV control. After the transductants expanded *in vitro* for several days, they were treated with biotin to label the PKC eta interactome, with or without 30 minutes of TCR stimulation, using T cells receiving no biotin for labeling as additional negative controls (**Figure 5C**). Mass spectrometry analysis of the biotinylated hits identified 3642 distinct proteins, 1183 of which passed basic cutoffs for expression in CD8^+^ T cells or enrichment over the mean signal of the negative controls (**Figure 5D**).

Applying stringent cutoffs for enrichment over negative controls (log_2_(FC) > 5 for anti-CD3 stimulated cells, log_2_(FC) > 3 for unstimulated; p_adj_ < 0.01 for both) identified candidate PKC eta- interacting proteins that could be categorized into those that interacted in resting T cells and those that interacted after TCR stimulation, including some that interacted in both resting and stimulated cells (27 proteins) (**Figure 5E**). Notably, while CARMA1 (*Card11*) interacted with PKC eta in unstimulated cells, BCL10 and MALT1 were not represented in the PKC eta interactome, in line with the observation that PKC eta poorly activates the CBM complex (**Figure 5A**). Proteins interacting with PKC eta in both stimulated and unstimulated conditions, particularly known scaffold proteins such as RLTPR/*Carmil2*^40,41^, are likely relevant to PKC eta function.

To globally assess biological roles of putative PKC eta “interactors”, we annotated these top 458 hits with their primary subcellular locations and assigned categories of cellular functions based on existing literature. We determined the percentage of hits in the unstimulated or stimulated cases per location or function and then computed the difference between the two stimulation cases (e.g. %localized to ER in stimulated - %localized to ER in unstimulated; **Figure 5F**). Analysis of the individual hits as well as the global trends in stimulation-dependent interaction partners revealed several main findings. First, PKC interacts with proteins whose primary localization is at many different subcellular sites, including the nucleus and endoplasmic reticulum, consistent with prior findings in COS-7 cells showing that PKC eta can localize to lipid membranes throughout the cell^33^. Second, PKC eta interacts with an abundance of negative regulators (including MAP4K1/HPK1, PTEN, RAB3GAP1, 5 members of the PP2A phosphatase complex^42^, and over 20 proteins involved in the ubiquitin/proteasome system). While these data do not specify the functional relationship between PKC eta and these interaction partners, they are suggestive of a model in which PKC eta functions by tuning the activities of a suite of negative regulators. Third, several protein complexes were statistically overrepresented among PKC eta’s interactome: along with PP2A, the OST N-glycosylation complex^43,44^ and EXOSC RNA degradation complex^45^ significant interacting complexes. Altogether, these data indicate that PKC eta is distinct from PKC theta in both its localization and regulatory roles: rather than promoting effector functions at the immune synapse, PKC eta interacts with protein complexes throughout the cell, connecting antigen signaling to a wide range of cell biology. These connections represent critical new avenues for considering how a kinase impacts cell fate choice in T_EX_ cells.

### Deletion of the PKC eta target CK1G2 improves the T_EX_ response to virus and cancer

Because of the association of PKC eta with T_EX-TERM_, we hypothesized that kinases downstream of PKC eta may broadly promote T cell exhaustion and dysfunction. This finding would open a new domain of signaling molecules that regulate T_EX_ cell fates and consequently a new class of targets for cancer immunotherapy.

Because T cell function could be improved in an *in vitro* exhaustion assay using the inhibitor D4476 (**Figure 4H**), which inhibits multiple forms of CK1, we tested the effects of individually deleting each of the CK1 genes that are expressed in CD8^+^ T cells during chronic infection (*Csnk1a1*, *Csnk1d*, *Csnk1g2*). P14 cells bearing these deletions, or a non-targeting *scr* guide, were adoptively transferred into mice, which were subsequently infected with LCMV Cl 13 (**Figure 6A**). P14 cells deleted for *Csnk1a1* showed a reduction in overall cell numbers, while deletion of *Csnk1d* caused no change in cell numbers, and most importantly, the cells deleted for *Csnk1g2* expanded significantly more than control cells by 8 days post-infection (**Figure 6B**). This finding suggested that deletion of *Csnk1g2* is highly beneficial for CD8^+^ T cells during chronic LCMV Cl 13 infection, and we focused our efforts on better understanding the effects of manipulating it.

**Figure 6.**
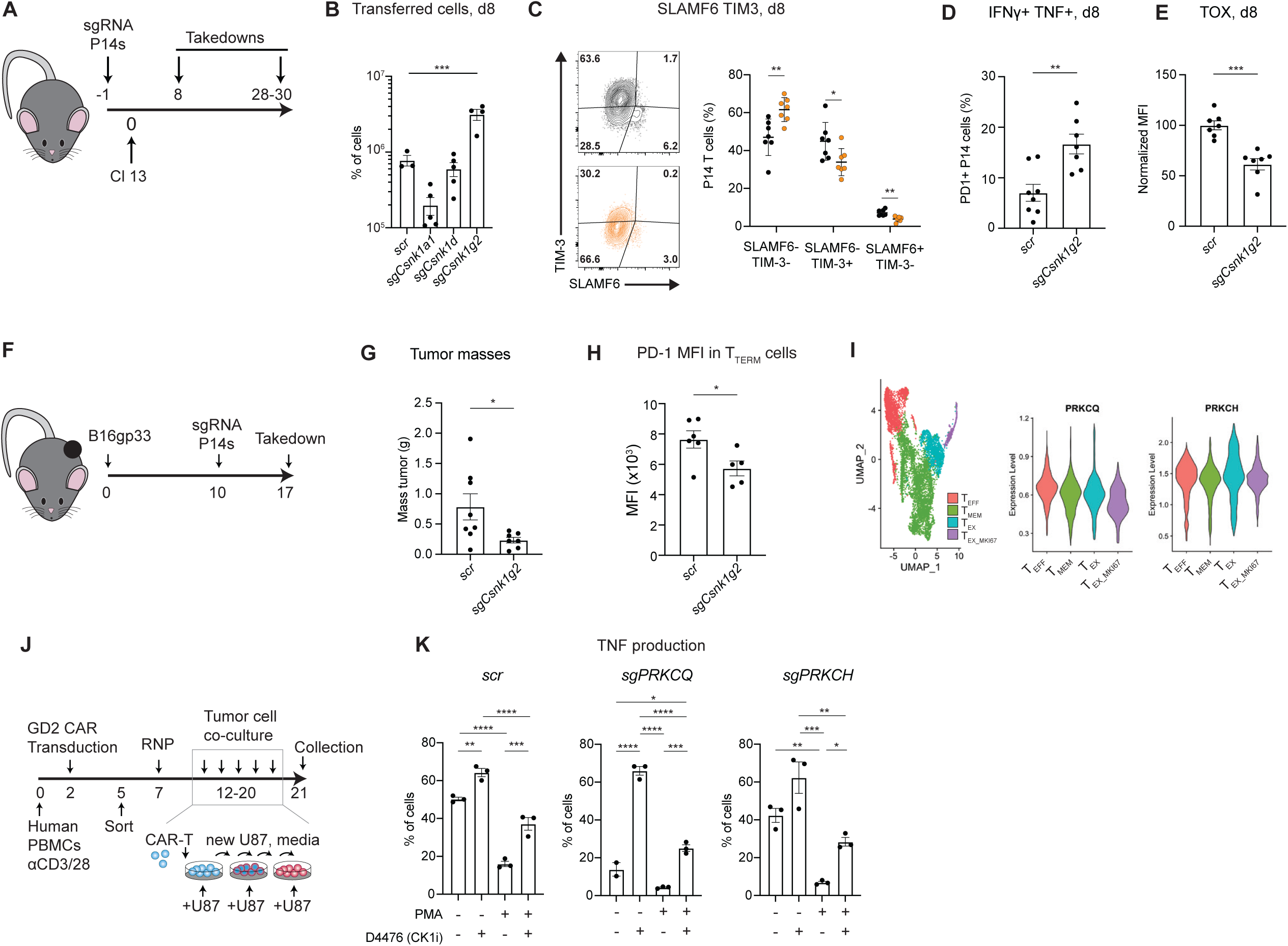
Deletion of the PKC eta target CK1G2 improves the T_EX_ response. (A) Timeline of casein kinase I gene deletions in LCMV Cl 13 adoptive transfer experiments. (B) Frequency of the indicated genotype transferred P14 T cells among all CD8^+^ T cells .(C-E) Representative data (C), frequencies (C-D), and MFIs (E) of markers of differentiation and functionality of P14 T cells of the indicated genotypes. (F) Timeline of *Csnk1g2* deletion in B16gp33 adoptive transfer experiments. (G) Masses of B16gp33 tumors at sacrifice timepoint by indicated genotype. (H) MFI of PD-1 in TTERM P14 T cells infiltrating B16gp33 tumors. (I) (Left) UMAP of integrated single cell RNA-seq datasets of CD8^+^ T cells isolated from human cancer biopsies (GSE146771, GSE99254, GSE98638), with cell subsets indicated by color. (Middle, right) Violin plots of *PRKCQ* and *PRKCH* expression by T cell subset. (J) Schematic of *in vitro* exhaustion experiments used to test the effects of CK1 inhibition and perturbations of PKC signaling on human CAR-T cells. (K) TNF production of *in vitro* exhausted human CAR-T cells by the indicated genotypes and treatments. Scatter plots show mean +/- s.e.m. Statistical significance was determined using Student’s t-test (panels C-E, G-H) or one way ANOVA (panels B, K), with significance shown as *P < 0.05, **P < 0.01, ***P < 0.001, ****P < 0.0001.

Compared to control *scr* cells at 8 d.p.i., P14 T cells deleted for *Csnk1g2* differentiate less frequently into SLAMF6^+^ TIM3^-^ T_EX-PROG_ or SLAMF6^-^ TIM3^+^. Instead, these cells preferentially differentiate into SLAMF6^-^ TIM3^-^ cells (**Figure 6C**) that produce significantly more IFNγ and TNF cytokines upon restimulation than *scr* cells and express less TOX (**Figure 6D-E**). These findings indicate that deletion of *Csnk1g2* causes T cells to exhibit greater effector function and become less exhausted during early timepoints of chronic infection, while remaining uncommitted to a terminal effector fate. Together, these data show that disruption of *Csnk1g2* in antigen-specific T cells leads to improvements in cell numbers and in effector function during the response to chronic viral infection.

We next tested whether deletion of *Csnk1g2* would also improve the anti-tumor function of CD8^+^ T cells. We implanted mice with B16-gp33 tumors and, 10 days post-implant, we adoptively transferred activated P14 T cells that had been deleted for *Csnk1g2* or a non-targeting control (**Figure 6F**). Indeed, mice that received P14 T cells lacking *Csnk1g2* had smaller tumors (**Figures 6G**), and the *sgCsnk1g2* P14 T cells expressed a reduced level of PD-1 (**Figure 6H**). Together, these data suggested that deletion of *sgCsnk1g2* enabled improved anti-tumor function of antigen-specific T cells.

### Human CAR-T cells require PKC theta for effector function and can be rescued by CK1 inhibition

To translate our work to human health, we first examined the expression of *PRKCQ* and *PRKCH* in an integrated analysis of scRNA-seq data of CD8 TILs from colorectal cancer^46^, non- small cell lung cancer^47^, and hepatocellular carcinoma samples^48^ (**Figure 6I**). Indeed, *PRKCH* expression increased significantly in T_EX_ cells isolated from these patients. To directly test the relevance of PKC and CK1 signaling to human cells, we conducted *in vitro* exhaustion experiments by co-culturing GD2 CAR-T cells with U87 target cells. Human CD8^+^ T cells were activated and purified from PBMCs and transduced with a lentiviral vector encoding a CAR specific for GD2. After sorting for CAR-T transductants, we electroporated the CAR-T cells with guide RNAs against *PRKCQ*, *PRKCH*, or a *scr* control. The CAR-T cells of each genotype were then subjected to *in vitro* exhaustion conditions that tested the effects of chronic PMA treatment or CK1 inhibition on their function and differentiation (**Figure 6J**). After 9 days of continuous exposure to antigen, we tested the effector function of CAR-T cells upon restimulation. Deletion of *PRKCQ* or co-culture with PMA caused major reductions in TNF production and a partial reduction of IFNγ (**Figure 6K**), consistent with our findings in mouse cells, while deletion of *PRKCH* had little effect on cytokine production. Meanwhile, treatment with the CK1 inhibitor D4476 boosted T cell functionality over its paired vehicle control samples. In particular, CK1 inhibition potently rescued TNF production in T cells lacking *PRKCQ*.

These findings suggest that our core findings on the PKC/CK1 signaling axis are conserved in human CAR-T cells. Altogether, our data suggest that PKC theta signaling is essential for effector function and maintenance of the antigen-specific T cell population in chronic antigen settings. In the absence of PKC theta, such as after activation-induced proteolytic degradation, the PKC eta/CK1 signaling axis promotes hyporesponsive, terminally exhausted cells. Our rescue experiments, via engineering of the non-degradable PKC theta K413R or inhibition of CK1, indicate that manipulation of distinct PKC signaling activities offers a rich set of therapeutic avenues in immuno-oncology.

## DISCUSSION

Here, we provide evidence that subsets of exhausted CD8 T cells respond differently to antigen by virtue of downstream differences in the signaling of two PKC paralogs, PKC theta and eta, with broad implications for the intersection of the cell biology of T cells and cancer immunotherapy.

We found that PKC theta is necessary for formation and maintenance of T_EX-PROG_ and for T cell effector functions, while PKC eta promotes T_EX-TERM_. PKC eta becomes dominant as T cells differentiate, as PKC theta is transcriptionally downregulated and simultaneously targeted for proteolytic degradation while actively signaling. Restoring PKC theta function by overexpressing a degradation-resistant variant of PKC theta, K413R, was sufficient to improve T cell function in cancer and chronic viral infection. PKC theta and eta execute these different genetic programs via different downstream signaling, which begins with PKC eta failing to activate the canonical PKC theta scaffold complex of CARMA1/BCL10/MALT1. Instead, PKC eta interacts with members of the OST glycosylation complex, the RNA degradosome, and the PP2A phosphatase, while promoting the activity of a distinct set of kinase targets. Indeed, inhibiting the putative PKC eta downstream target CK1G2 rescued T_EX_ function *in vivo* and in human CAR-T cells *in vitro*.

In general, the different responses of PKC theta and PKC eta to antigen signals may reflect the different biology of T_EX-PROG_ and T_EX-TERM_, respectively. Our work here and other studies^36,40,41^ show that PKC theta and CARMA1 signaling favors pathways that enable proliferation and effector function. Meanwhile, PKC eta’s interaction partners reflect a partial maintenance of effector function^49^ but a potential switch to more housekeeping or regulatory functions. Considering these findings together with the observation that PKC theta is expressed highly in T_EX-PROG_ and T_EX-EFF_, we propose that the loss of PKC theta is a driving feature of the loss of functionality in T_EX-TERM_, and that maintaining a high level of PKC theta activity is a key component in both the plasticity of T_EX-PROG_ and skewing T cells towards the more functional subset of T_EX-EFF_. Though we have identified differences in the pathways downstream of PKC theta and eta, these findings raise new questions about the implications and mechanisms of these differences. Phosphoproteomics and follow-up genetics indicated that CK1G2 is a downstream target of PKC eta and that it has a detrimental effect on CD8 T cell function. However, how PKC eta activates it or how CK1G2 negatively impacts T cell function is currently unclear. Beyond kinase cascades, our proximity ligation data also suggest a role for PKC eta in regulating diverse biological processes such as glycosylation and mRNA degradation. Future studies will investigate the roles of these interactions in T_EX-TERM_ cells.

Despite PKC theta and eta having similar specificities for model peptides^29^, these kinases drive distinct cell biology at different subcellular locations. PKC theta signals through CARMA1 at the immunological synapse between a T cell and an antigen presenting cell^50^. By contrast, we found that PKC eta interacts not only with plasma membrane proteins, but upon antigen stimulation, interacts with proteins that localize to the ER and nucleus. Indeed, prior studies have shown that PKC eta can localize to multiple sites^50^, including the ER and nucleus, dependent upon its pseudosubstrate and C1 domain^33^, one of several studies to indicate that PKCs C1 and C2 domains for regulatory roles aside from recognition of activating ligands^22,34,51,52^.

Proximity ligation also suggests an interaction between PKC eta and the PP2A phosphatase complex. This interaction could reflect PKC eta’s interaction with its own negative regulator^53^ but is also suggestive of a role for PKC eta in regulating PP2A^54^, whose targets include Akt, mTOR, MEK^42^ and whose subunit *Ppp2r2d* was the lead hit in a genome-wide screen for T cell anti-tumor function^55^. Broadly, PKC eta interacts with negative regulators beyond PP2A: *Map4k1*, *Rab3gap1*, *Pten* all associate with PKC eta in both stimulated and unstimulated conditions. In concert with findings that PKC eta is essential for CTLA-4 signaling in regulatory T cells^32^, these data suggest that PKC eta has widespread connections to negative signaling regulators in T cells and are in line with recent work on PKC as a tumor suppressor^53,56,57^. Indeed, phosphoproteomics presented here and elsewhere may reflect not only direct downstream targets of a kinase, but also the effects of phosphatases under the kinase’s control.

Our study presents both immediate applications and new concepts in cancer immunotherapy. PKC theta and PKC eta promote T_EX-PROG_ and T_EX-TERM_ respectively, and we showed that expression of a degradation-resistant PKC theta or ablation of the putative PKC eta target CK1G2 could improve anti-tumor T cell function. Beyond the therapeutic applications of these two approaches, their successes illustrate the potential of our strategy to identify and exploit subset-specific signaling differences in the design of new immunotherapies. Prior studies have highlighted the potential of engineering CAR-T cells by overexpressing key transcription factors^8,58,59^, or have amplified proximal TCR signaling by small molecule approaches^60^, or by blockade of inhibitory receptors^61^. Our work highlights the potential of manipulating downstream elements in signaling cascades which bestow uniquely beneficial or detrimental qualities to the differentiation states that employ them.

In total, our results illustrate that subsets of exhausted T cells respond differently to antigen via different signaling cascades downstream of PKC theta and eta. These findings indicate that understanding how core signaling pathways vary by differentiation state holds promise, and perhaps is essential, for the development of future immunotherapies.

## Materials and Methods

Key Resources Table

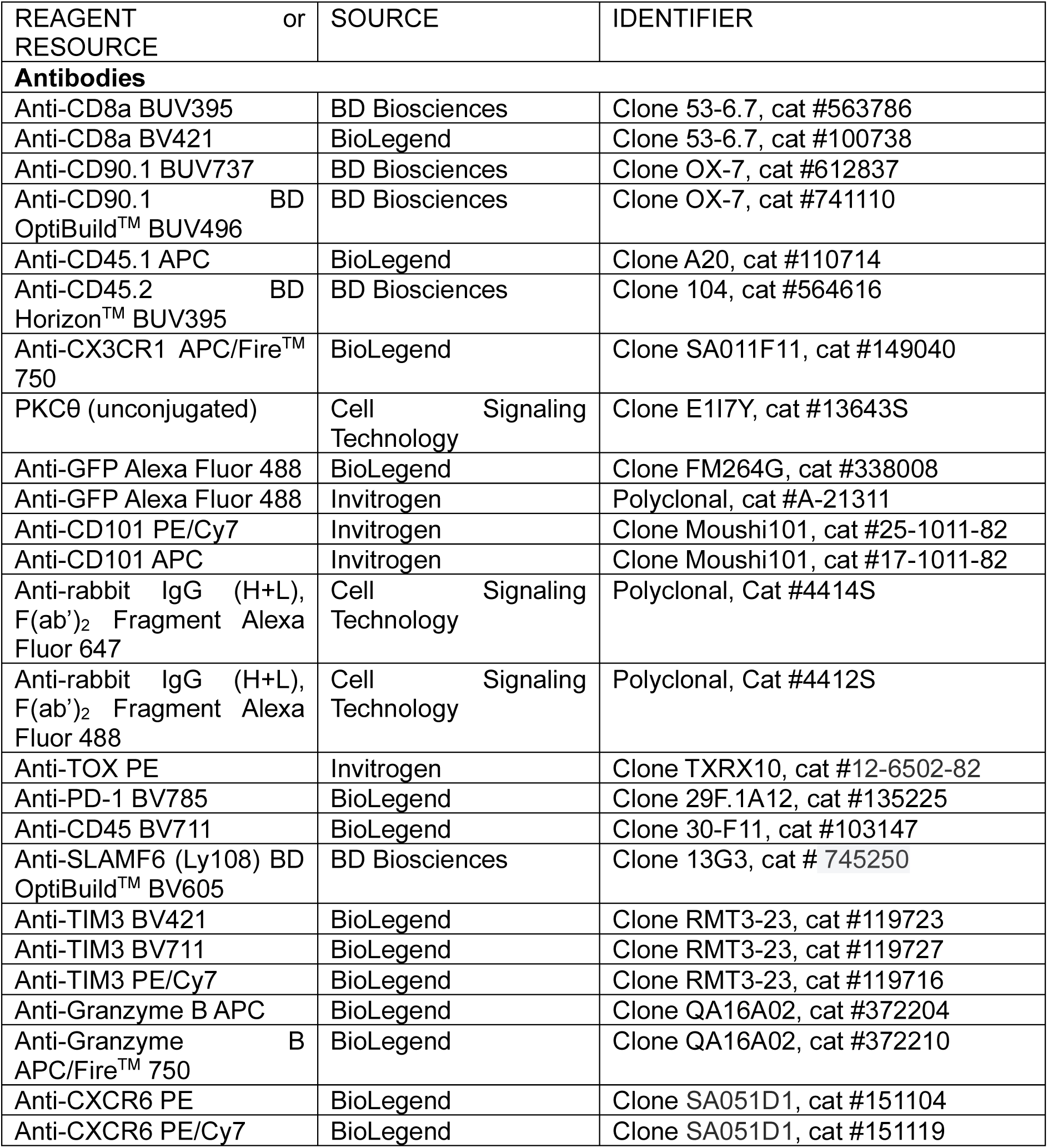
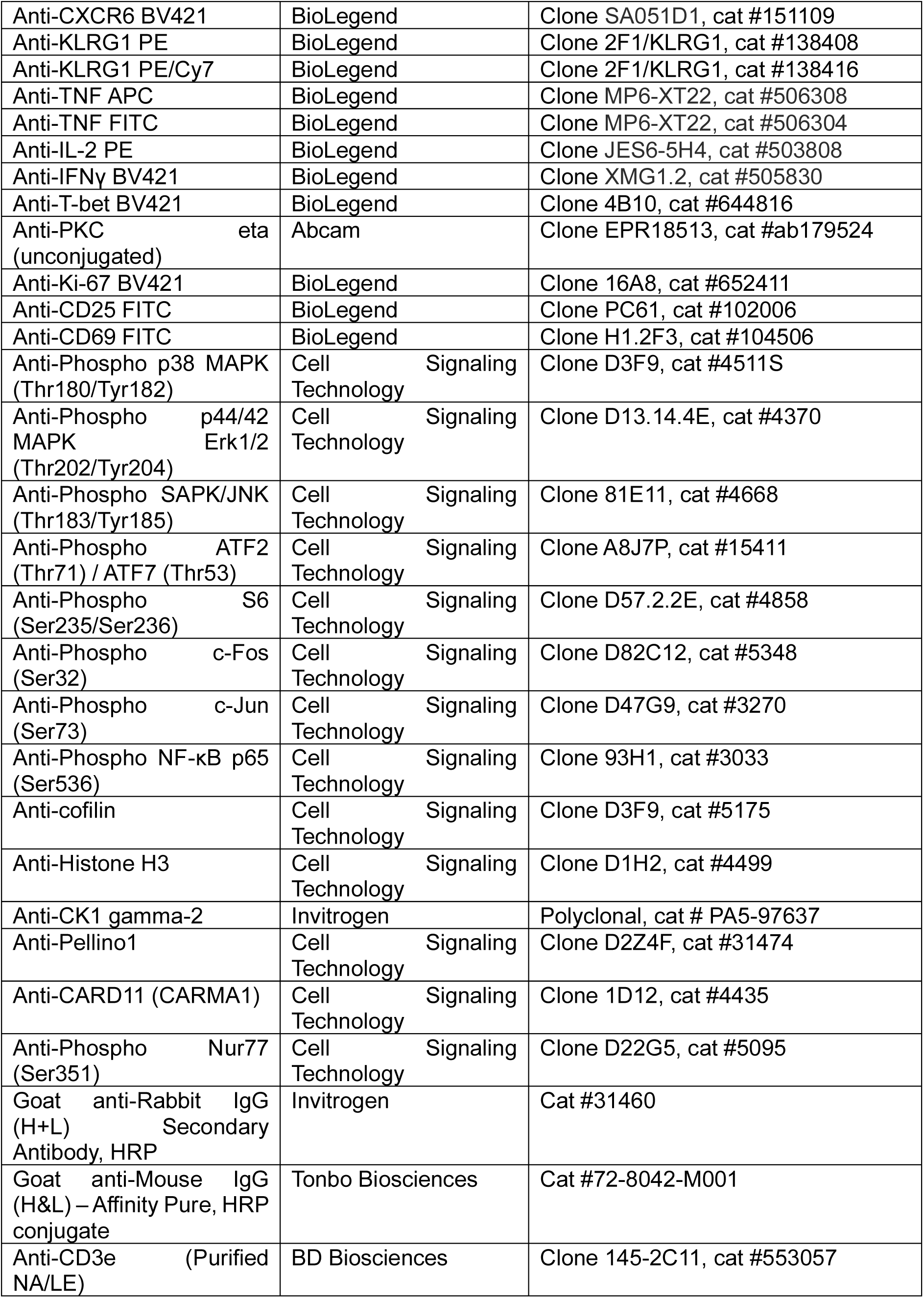
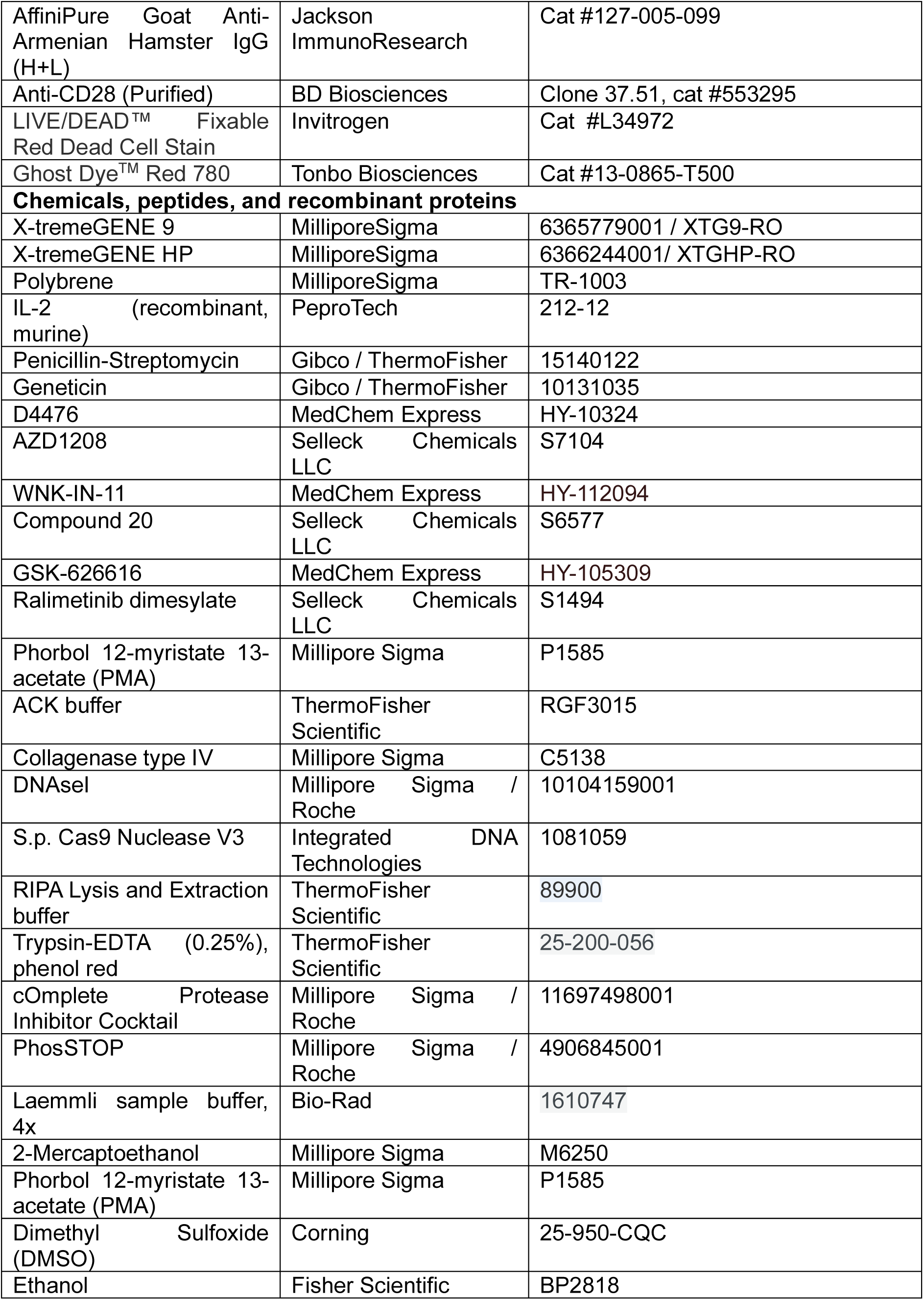
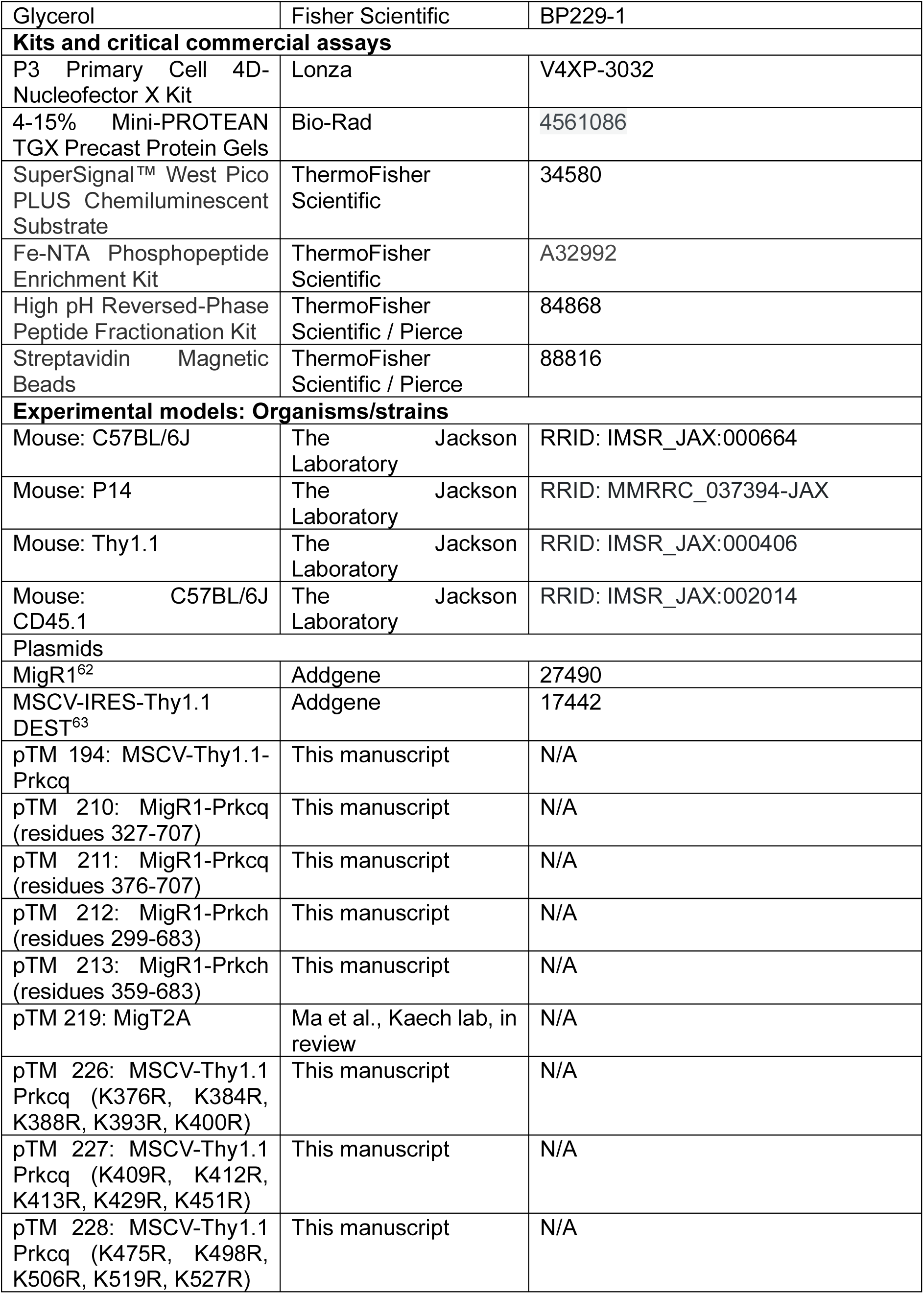
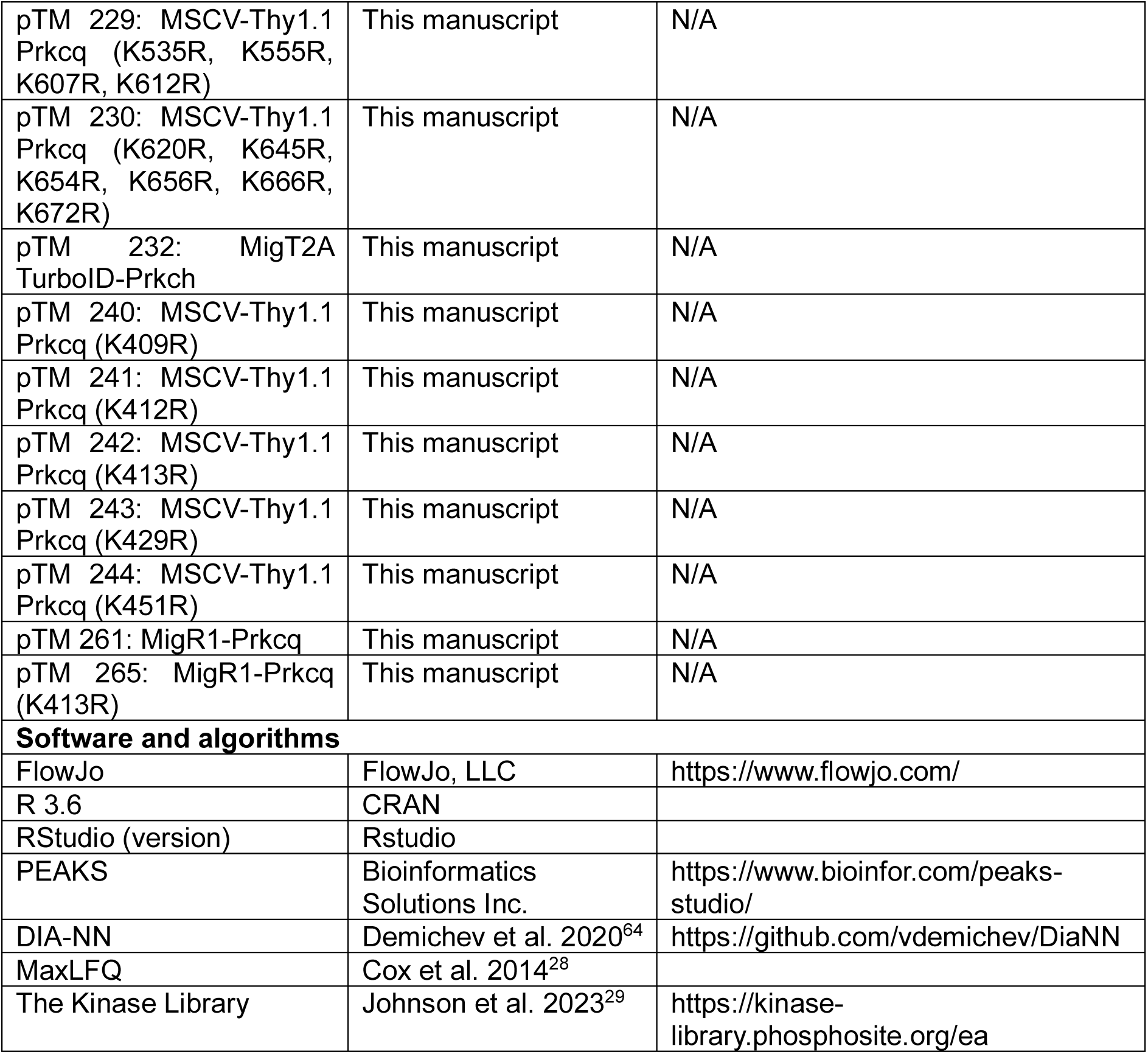

### RESOURCE AVAILABILITY

#### Lead contact

Further information and requests for resources and reagents should be directed to and will be fulfilled by the lead contact, Susan M. Kaech.

#### Materials availability

All plasmids newly created for this work are available upon request to the corresponding author.

#### Data and code availability

All proteomic data will be uploaded to the PRIDE server. Code for analysis of phosphoproteomics and TurboID data will be available upon request.

### EXPERIMENTAL MODEL AND SUBJECT DETAILS

#### Mice

C57BL/6 (B6) mice were used for all experiments. C57BL/6 mice were purchased from Jackson Laboratory. For adoptive transfer experiments, T cells were collected from TCR transgenic P14 C57BL/6 mice (TCR specific for LCMV D^b^GP_33-41_) mice expressing congenic markers (Thy1.1 or Ly5.1). The donor cells were transferred into sex- and age-matched recipients. P14 mice were backcrossed to Jackson Laboratory C57BL/6 mice three times every two years during the study to avoid genetic drift-based cell rejection. For in vitro exhaustion experiments, T cells were collected from P14+ mice and activated with GP_33-41_ peptide *in vitro*. Both male and female mice were used unless stated otherwise. All mice were used according to Institutional Animal Care and Use Committee (IACUC) guidelines at Salk Institute for Biological Studies.

### METHOD DETAILS

#### Cell culture media and buffers

The names and recipes for all the buffers listed in subsequent methods sections are described here. FACS buffer: PBS supplemented with 2% FBS and 0.02% sodium azide. T cell media: RPMI 1640, 10% FBS, penicillin/streptomycin (Gibco), 2 mM L-glutamine, 50 μM beta-mercaptoethanol. Β16-gp33 media: DMEM, 10% FBS, penicillin/streptomycin (Gibco; final concentrations of 100 units/mL penicillin and 100 μg/mL streptomycin), and 100 μg/mL geneticin to maintain expression of gp33. Tumor digestion media: DMEM, 10% FBS, penicillin/streptomycin, 0.5 mg/mL collagenase type IV (Millipore Sigma), 0.1 mg/mL DNAseI.

#### LCMV Cl 13

LCMV Cl 13 was produced according to prior studies^65^. Mice 4-8 weeks old were infected intravenously (i.v.) with 2×10^6^ pfu Cl 13. Adoptive transfers were performed as described in the adoptive transfer section, and tissues were collected from euthanized mice and processed as described in T cell isolation. Blood was collected via retro-orbital bleeding of isoflurane-anesthetized mice. All *in vivo* studies were performed according to guidelines approved by the Salk Institute Institutional Animal Care and Use Committee (IACUC).

#### Subcutaneous tumor models

Subcutaneous tumor models were produced by implanting B6 mice 4-8 weeks old on their flanks with 4×10^5^ B16-gp33 cells. Mice were shaved at the implant site prior to injection. B16-gp33 cells were expanded and collected, at low passage number, by treating with trypsin for 3 minutes. Cells were counted, rinsed, and resuspended at a concentration of 4×10^6^ cells/mL in PBS for injection. Tumor measurements were performed with calipers starting at d7 post-implantation and were performed at least twice per week until endpoint. Tumor volumes were calculated from caliper measurements assuming a prolate ellipsoid shape (V = 0.5233 x length^2^ x width), where the length is the longer axis of the caliper measurements. Immediately prior to sacrifice, one final caliper measurement of tumor size was taken. All *in vivo* studies were performed according to guidelines approved by the Salk Institute IACUC.

#### T cell isolation from spleen or tumor

##### T cells from spleen

Spleens were collected from euthanized animals and immediately placed in T cell media on ice. Spleens were gently mashed on a 70 μm nylon filters using a 1 mL syringe, and the filters were washed generously with T cell media. Splenocytes were spun down at 500 rcf, media was removed, and red blood cells were lysed by incubating the splenocytes in 1 mL ACK buffer for 4 min at room temperature. Lysis was quenched by addition of 5 mL T cell media. Cells were resuspended in T cell media and prepared for experimental use.

##### T cells from tumor

Subcutaneous tumors were dissected from mice, weighed, diced with razor blades in a 60 mm petri dish, and placed on ice. The tumor slurry was suspended in 5-10 mL of tumor media and shaken at 200 rpm in a shaker incubator at 37 °C for 30 min. The slurry was then mashed through a 70 μm nylon filter using a 1 mL syringe plunger to produce a single cell suspension. The filter was then rinsed with extra tumor media to collect all the cells. The cell suspension was spun at 450 rcf for 3 min at 4 °C, the supernatant was poured off, and the pellet was treated with 2 mL ACK buffer for 5 min at room temperature to lyse red blood cells. The ACK lysis was quenched with 10 mL tumor media, and cells were counted and then used for further experiments. All *in vivo* studies were performed according to guidelines approved by the Salk Institute IACUC.

#### Flow cytometry and antibodies

Following isolation of cells, single cell suspensions were stained with anti-CD16/32 (BioLegend) on ice for 15 min to block Fc receptors. Antibody cocktails for surface proteins were prepared up to 1 h in advance. Antibodies were added to FACS buffer, mixed well, and kept on ice. Immediately before staining cells, LIVE/DEAD Fixable viability stain was added to the antibody cocktail, mixed well, and cells were stained. For stains of 3×10^6^ cells or fewer, cells were stained in 50 μL of Ab cocktail; for stains greater than 3×10^6^ cells, the staining volume was increased to 100 μL. Surface and viability stains were done for 30 min on ice. Cells were rinsed once in FACS buffer. Centrifugation to collect cells and exchange incubation media was done at 450 rcf for 3 min at 4 °C for unfixed cells; after fixation, described below, centrifugation was done at 550 rcf, again for 3 min at 4 °C.

To measure cytokine production, cells were resuspended in 200 μL T cell media and stimulated. For LCMV experiments, P14 T cells were stimulated with 100 ng/mL LCMV gp33 (KAVYNFATC) in the presence of 5 μg/mL Brefeldin A (BioLegend 420601) for 5 h at 37 °C; for tumor experiments, T cells were instead stimulated with 10 ng/mL PMA and 1 μg/mL ionomycin for 4 h.

For intracellular staining of cytokines, cells were fixed for 1 min at room temperature with BioLegend Fixation Buffer (PBS with 4% paraformaldehyde, catalog number 420801). Cells were spun down, the fixation buffer was removed, and then cells were incubated for 30 min on ice in FoxP3 Transcription Factor Fixation/Permeabilization buffer. For intracellular stains of panels that did not include cytokines, the 1 min fixation step was omitted. Following fix/perm, cells were rinsed once in eBioscience Permeabilization buffer, spun down, and then stained with antibodies against other intracellular proteins. The intracellular Ab cocktail was made in Permeabilization buffer and cells were stained for 45 min on ice. Cells were rinsed twice with Permeabilization buffer, and then if required (such as for PKC theta or eta), stained with a fluorescently conjugated secondary antibody in Permeabilization buffer for 30 min on ice. After secondary Ab staining, cells were rinsed two more times in Permeabilization buffer. Then when staining was complete, cells were resuspended in FACS buffer and prepared for flow cytometric analysis. Samples were processed on an LSR II or A3 flow cytometer (BD Biosciences) and data were analyzed with FlowJo v10.

#### *In vitro* T cell exhaustion assays

In vitro T cell exhaustion assays were performed by continuously co-culturing P14 T cells with B16-gp33 cells. This protocol is conceptually similar to previously published protocols^20,21^. The timeline of the assay is referred to relative to days post-activation for the T cells.

*Day -2*. B16-gp33 cells were started fresh from liquid nitrogen storage or split at low passage number into a new flask containing B16 media.

*Day 0*. P14 T cells were activated with 100 ng/mL gp33 and 10 ng/mL recombinant murine IL-2. B16-gp33 cells were split to have one T175 flask.

*Day 1*. if genetic modification of T cells was required, it was performed following the protocols described below in the CRISPR/Cas9 RNP electroporation or retroviral transduction sections. Regardless of whether genetic modification was performed or not, T cells were exchanged into fresh T cell media.

*Day 2*. B16-gp33 cells were removed from the flask by treating them with Versene for 3 min at 37 C. The cells were exchanged into B16-gp33 media and then irradiated to induce cell cycle arrest to facilitate the maintenance of the assay over the course of a week. A dose of 45 Gray was administered by placing cells in a conical tube 25 cm from a Co-60 source for the required time (5-8 min, as the source decayed over the course of this study). Cells remained over 95% viable after irradiation. Cells were plated at 5×10^4^ cells per well of a 96 well flatbottom plate, 1.5×10^6^ cells per well of a 6 well plate, or otherwise scaled appropriately by vessel surface area.

*Day 3*. The splenocyte mixture containing activated P14 cells was collected, and cells were counted. 2.5×10^4^ splenocytes were plated in each well, a ratio of 1:2 of splenocytes to B16-gp33 cells, roughly corresponding to 1:5 P14 to B16-gp33 cells. The B16-gp33 media, containing geneticin, was removed, and wells were rinsed with T cell media prior to addition of P14 cells in T cell media. The T cell / B16-gp33 co-culture was supplemented with cytokines or drugs as described in the main text, and the total culture volume was 200 μL in each well of a 96 well plate. Our standard conditions for enforcing the most severe level of exhaustion entailed using IL-2 at 10 ng/mL and PMA at 1 μg/mL (diluted 10,000x from a 10 mg/mL stock in ethanol).

*Days 4-6*. Nothing was done to the culture on day 4. On day 5, T cells were collected from the plate using a multichannel pipette by thoroughly resuspending media in each well of the plate, plated in a roundbottom plate to centrifuge the cells, and T cells were added with fresh media, cytokines, and drugs as appropriate to new wells of B16-gp33 cells. Again, B16-gp33 containing wells were rinsed with T cell media prior to addition of T cells. On day 6, this media exchange procedure was repeated.

*Day 7*. T cells were collected with pipetting. Versene was used where necessary if T cells remained bound to surviving B16-gp33 cells in certain conditions: using Versene rather than trypsin avoids potential cleavage of surface antigens that may be stained in flow cytometry. T cells were then used for downstream assays including flow cytometry and gene expression analysis.

#### Chemical biology and drug treatments

*Inhibitor screening of kinases downstream of PKC theta or eta*: GSK-626616 (MedChem Express) was used to inhibit DYRK^66^ and was diluted from a 10 mM stock to a final concentration of 1 μM. AZD1208 (Selleck Chemicals) was used to inhibit PIM kinases^67^ was diluted from a 10 mM stock in DMSO to a final concentration of 1 μM. and inhibitor. Compound 20 (Selleck Chemicals) was used to selectively inhibit PKC theta^68^ and was diluted from a 50 mM stock in DMSO to a final concentration of 5 μM. Ralimetinib dimesylate (Selleck Chemicals) was used to inhibit p38^69^ and was diluted from a 10 mM stock in DMSO to a final concentration of 5 μM. WNK-IN-11 was used as a WNK inhibitor^70^ and was diluted from a 10 mM stock in DMSO to a final concentration of 1 μM. D4476 was used as a pan-CK1 inhibitor^71^ and was diluted from a 10 mM stock in DMSO to a final concentration of 10 μM. *PKC theta degradation assays*: To induce PKC theta degradation and/or to measure the effects on T cells of chronic PKC agonism, cells were continuously cultured in phorbol 12-myristate 13-acetate (PMA). For Nanostring gene expression analysis, T cells were incubated with PMA at a concentration of 1 μg/mL in T cell media (with 10 ng/mL IL-2) from days 3 to 7 post-activation, changing media every day. For degradation timecourse assays and PKC theta mutant screening, the same culture and PMA conditions were used, but for up to a maximum of 24 hours of PMA treatment.

#### Nanostring gene expression analysis

Nanostring gene expression analysis was performed on cells that were subjected to the *in vitro* exhaustion protocol described in the above section, with minor changes. The cells for Nanostring analysis did not receive antigen beyond initial peptide-activation, but they did receive media changes, including PMA or a DMSO vehicle control, on the same schedule as the *in vitro* exhaustion protocol (new media, IL-2, and drugs on days 3, 5, 6, and 7, with sample collection on day 8 for this analysis). Two technical replicate wells of a flatbottom 96 well plate were pooled, and 3×10^4^-5×10^4^ cells were collected for RNA isolation (RNA Clean & Concentrator-5, Zymo # R1013). The isolated RNA was hybridized with a nCounter custom probe set (108 probes) according to the manufacturer’s protocol. The hybrid RNA samples were loaded onto a Nanostring nCounter MAX Profiler. The expression data was normalized to expression of a panel of positive control housekeeping genes (*Cd8a*, *Cd2*, *B2m*) and internal negative control probes. Phenotypes of the cells were validated in parallel by performing flow cytometry on cells that were collected from the same samples but not used for Nanostring.

#### Plasmid cloning and molecular biology

Retroviral overexpression plasmids were created using 2x NEBuilder HiFi DNA Assembly master mix (New England Biolabs #2621) to perform Gibson assembly. Genes of interest were inserted into several different backbones (MigR1^62^, Addgene #27490; MSCV-IRES-Thy1.1 DEST^63^, Addgene #17442). MigR1 was digested with XhoI and MSCV-IRES-Thy1.1 was digested with NotI to linearize the vectors before Gibson assembly. A Kozak sequence (GCCACC) was added before the start codon of inserted genes to facilitate translation. Long (>300 bp) DNA inserts were synthesized at Twist Bioscience and oligonucleotides were synthesized at Integrated DNA Technologies.

For non-degradable *Prkcq* mutant screening, the WT *Prkcq* gene sequence was divided into 4 fragments, and WT and mutant versions of each fragment (containing Lys to Arg mutations in the region) were synthesized to facilitate mutant scanning in blocks of 4-6 Lys residues (e.g. Frag 1 (WT), Frag 2 (containing muts K409R, K412R, K413R, K429R, K451R) Frag 3 (WT), Frag 4 (WT)). This mixing and matching of modules allows for rapid generation of many gene variants using a handful of short, directly synthesized DNA segments.

#### CRISPR/Cas9 deletion of genes in primary T cells

Following isolation of splenocytes, CD8 T cells were purified by negative selection. A negative selection cocktail was prepared by incubating streptavidin-coated magnetic beads with biotin conjugated Abs against CD4, MHC-II, CD11c, CD11b, Ter119, B220, CD49b, and TCR γ/δ. For CRISPR/Cas9 RNP deletion of genes encoding the PKCs (*Prkcq* or *Prkch*, as well as the scramble sgRNA negative control used in the PKC experiments), the RNP electroporation was performed 24 h post-activation of T cells so that deletion of *Prkcq* would not interfere in the priming of naïve T cells. Otherwise, RNP electroporation was performed on naïve cells. Unless otherwise stated, all guide sequences were obtained from the Brie library^72^, and their deletion of the target gene was validated by flow and/or Western blot.

RNP complexes were formed by incubating 36 pmol Cas9 (IDT) with 300 pmol sgRNA (Synthego) in 5 mL RNAse-free water for 10 min at room temperature. As the RNP complexes formed, 2×10^6^ purified CD8 T cells were aliquoted into a 1.7 mL tube for each electroporation reaction, and cells were rinsed in PBS. When cells were ready for electroporation, they were centrifuged, and the supernatant was completely removed. Cells were electroporated using the Lonza P3 Primary Cell 4D-Nucleofector X kit according to manufacturer instructions and as described previously^15^. Briefly, the cell pellet was resuspended in 20 μL of a mixture of P3 buffer and Supplement 1, and the cell suspension was immediately transferred to a tube containing the RNP complex mixture. This mixture was again rapidly mixed via pipetting and transferred to a cuvette for electroporation using the Lonza program DN100. Following electroporation cells were removed from the cuvette and allowed to recover in 5 mL of T cell media pre-warmed to 37 °C. After 15 min of recovery at 37 °C, cells were pelleted and prepared for experimental use.

#### Retroviral transduction of primary T cells

T cells were engineered to overexpress target genes by retroviral transduction. Ecotropic retrovirus was packaged in HEK 293T cells by transfecting the HEK cells with a target plasmid of interest (described in plasmid cloning above) and pCL-Eco (Addgene #12371). The conditions listed here describe one transfection reaction worth of material, and this recipe was scaled accordingly to the amount of retrovirus needed to transduce sufficient T cells.

The day before transfection, 4×10^5^ HEK cells were plated in 2 mL HEK media (DMEM, 10% FBS, penicillin/streptomycin) in one well of a 6 well plate; each well of a 6 well plate provided enough virus to transduce 3×10^6^ splenocytes from a P14 mouse or 7.5×10^5^ purified CD8 T cells. On transfection day, each transfection reaction was made in a 1.7 mL tube and contained 1 μg of target plasmid, 0.5 μg pCL-Eco, and 6 μL X-tremeGENE 9 (MilliporeSigma), and a volume of OptiMEM (ThermoFisher 31985062) to reach 100 μL (X-tremeGENE added last); X-tremeGENE HP (MilliporeSigma) was also used to improve transfection efficiency. The reaction was mixed by gently flicking the tube. After 15 min incubation at room temperature, the mixture was added dropwise to the HEK cells and the plate was gently swirled and then incubated at 37 °C overnight. About 18 h post-transfection, the HEK media was removed and replaced with 2 mL fresh HEK media, gently adding media along the side of the well to avoid disrupting the HEK cells.

The same day of the HEK media replacement, splenocytes were isolated as above and T cells were activated. For experiments using P14 T cells, the leukocytes remaining after red blood cell lysis were resuspended in 10 mL T cell media, gp33 peptide was added to a final concentration of 100 ng/mL, and recombinant murine IL-2 (PeproTech 212-12) was added to a final concentration of 10 ng/mL. For experiments not using P14 T cells, CD8 T cells were isolated via negative selection as described above, resuspended to 3×10^6^ cells/mL in T cell media (with 10 ng/mL IL-2), and activated via platebound anti-CD3/28 stimulation. Once T cells were placed in their activation media, they were incubated at 37 °C overnight.

The following day, 18-24 hours post-activation, the retrovirus containing supernatant was collected from HEK 293T cells and centrifuged at 300 rcf at room temperature for 10 minutes to remove any stray HEK cells. For each transduction, T cells were counted and adjusted to plate 3×10^6^ activated P14 splenocytes or 7.5×10^5^ purified CD8 T cells per well in a 12 well plate, and the retrovirus from one transfection was added to each well of T cells. Polybrene (MilliporeSigma

TR-1003) was added to a final concentration of 9 μg/mL and gently mixed. T cells were transduced by centrifuging the transduction mixtures at 1500 rcf for 90 minutes at 32 °C; the brake speed of the centrifuge was reduced to avoid disturbing nascent retroviral infections. Following the spin, the transduction plate was gently removed from the centrifuge and incubated at 37 °C for 3 h to allow infection to complete. After this 3 h rest, transductions were spun again at 600 rcf at 32 °C for 3 min to adhere T cells to the plate, and the media was gently removed. The transduced T cells were then either given fresh T cell media with 10 ng/mL IL-2 if they were to be used for *in vitro* experiments, or they were prepared for adoptive transfer for *in vivo* experiments. The transduction protocol used here is based on prior work^73^.

#### Adoptive T cell transfer

To prepare T cells for adoptive transfer, the cells were rinsed once in PBS and then resuspended in PBS. T cells were always transferred via retro-orbital injection of 100 μL per mouse. The activation state of the T cells, the timing of transfer, and the number of cells were adjusted accordingly for each immune challenge as follows.

For all LCMV Cl 13 experiments, 1.5×10^4^ purified T cells or 7.5×10^4^ transduced P14 splenocytes were adoptively transferred. T cells that were either transduced or were electroporated 24 h post-activation were transferred into mice that had also been infected 24 h prior, so as to match the activation timeline for the transferred and endogenous T cells. For T cells that were electroporated while naïve, they were transferred into mice 24 h before the mice were infected.

For subcutaneous tumor experiments, T cells were transferred on day 10 post-implantation, when tumors had become palpable. Unless otherwise stated, 2×10^5^ purified T cells (or 1×10^6^ P14 splenocytes) were transferred 24 h post-activation.

#### Proteomics: phosphoproteomics and TurboID

##### Cell culture for phosphoproteomics

CD8 T cells were negatively selected and pooled from 5 animals, activated overnight by platebound anti-CD3/28 stimulation, and subsequently electroporated with RNP complexes containing guides against *Prkcq*, *Prkch*, or a scrambled non-targeting control. Cells were expanded *in vitro* until 6 days post-activation and then they were prepared for re-stimulation for phosphoproteomic analysis. Prior to sample collection, 5×10^7^ cells per genetic background were aliquoted per re-stimulation condition. Cells were plated on 10 cm dishes with or without anti-CD3 coating. Cells were re-stimulated for 30 min and then immediately placed on ice and cells were collected. The cell pellets were spun down, the supernatant was removed, the pellets were snap frozen in liquid nitrogen.

##### Sample preparation for mass spectrometry

Samples were lyophilized overnight and reconstituted in 200 μl of 6M Guanidine-HCl per 1 mg of protein. The samples were then boiled for 10 minutes followed by 5 minutes cooling at room temperature. The boiling and cooling cycle was repeated a total of 3 cycles. The proteins were precipitated with addition of methanol to a final volume of 90% followed by vortexing and centrifugation at maximum speed on a benchtop microfuge (14000 rpm) for 10 minutes. The soluble fraction was removed by flipping the tube onto an absorbent surface and tapping to remove any liquid. The pellet was resuspended in 200 μl of 8 M urea made in 100 mM Tris-Cl pH 8.0. TCEP was added to final concentration of 10 mM and chloro-acetamide solution was added to final concentration of 40 mM and vortexed for 5 minutes. The solution was then acidified using TFA (0.5% TFA final concentration). 3 volumes of 50mM Tris pH 8.0 were added to the sample to reduce the final urea concentration to 2 M. Trypsin was in 1:50 ratio of trypsin and incubated at 37°C for 12 hours. The solution was once again acidified using TFA (0.5% TFA final concentration) and mixed. Samples were desalted using Thermo C18-Stage Tips (cat #87782 and #87784) and 100 mg C18-solid phase extraction (Waters cat # WAT023590) as described by the manufacturer protocol. The eluted peptides are dried in a speed vac. The peptide concentration of sample was measured using BCA.

To prepare proteins for global phosphoproteomics, phospho-peptides were enriched using High-Select Fe-NTA Phosphopeptide Enrichment (A32992 Thermo Scientific). The enriched phosphopeptide fraction was then further fractionated using Pierce™ High pH Reversed-Phase Peptide Fractionation Kit (Pierce™ High pH Reversed-Phase Peptide Fractionation Kit Catalog number: 84868). Fractionation protocol as described by the manufacturer kit. Eight fractions were generated from this step.

To prepare peptides for TurboID analysis, biotinylated peptides were pulled down using streptavidin magnetic beads. To prepare the magnetic beads, 50 μL (0.5 mg) of Pierce Streptavidin Magnetic Beads (Pierce cat #88816) were aliquoted into a 1.5 mL microcentrifuge tube. The beads were separated on a magnetic stand and the supernatant was discarded. 1 mL of PBS was added to the tube, and the tube was inverted and vortexed gently to mix. The beads were collected with a magnetic stand, and the supernatant was discarded again. The beads should never dry during this process and can be stored in PBS if needed. The lyophilized sample of biotinylated and digested peptide samples were dissolved in 1 ml PBS, and this mixture was added to beads and incubated with mixing at room temperature for 1-2 hours. Beads were washed three times by adding 1 mL of PBS with 1 mL of 5% acetonitrile in PBS, and then a last wash was performed in ultrapure water. Excess liquid was completely removed from the beads using a micropipette, and biotinylated peptides were eluted by adding 0.3 mL of solution containing 0.2% TFA, 0.1% formic acid, and 80% acetonitrile in water. The beads were centrifuged at 1000 g and the first elution of biotinylated peptides was transferred to an Eppendorf tube. A second elution of 0.3 mL was boiled for 5 min for maximum release of peptides from the beads. A third elution of 0.3 mL was boiled for 5 min for maximum release of peptides from the beads. The eluted samples were dried in a speed vac.

##### LC-MS/MS analysis

For TurboID analysis, trypsin-digested peptides were analyzed by ultra high pressure liquid chromatography (UPLC) coupled with tandem mass spectroscopy (LC-MS/MS) using nano-spray ionization. The nanospray ionization experiments were performed using a TimsTOF 2 pro hybrid mass spectrometer (Bruker) interfaced with nano-scale reversed-phase UPLC (EVOSEP ONE). Evosep method of 30 SPD (samples per day) was utilized using a 10 cm × 150 μm reverse-phase column packed with 1.5 μm C18-beads (PepSep, Bruker) at 58°C. The analytical columns were connected with a fused silica ID emitter (10 μm ID; Bruker Daltonics) inside a nanoelectrospray ion source (Captive spray source; Bruker). The mobile phases comprised 0.1% formic acid (FA) as solution A and 0.1% FA/99.9% acetonitrile as solution B. The mass spectrometry setting for the TimsTOF Pro 2 are as following: PASEF method for standard proteomics. The values for mobility-dependent collision energy ramping were set to 95 eV at an inversed reduced mobility (1/*k*_0_) of 1.6 V s/cm^2^ and 23 eV at 0.73 V s/cm^2^. Collision energies were linearly interpolated between these two 1/*k*_0_ values and kept constant above or below. No merging of TIMS scans was performed. Target intensity per individual PASEF precursor was set to 20 000. The scan range was set between 0.6 and 1.6 V s/cm^2^ with a ramp time of 166 ms. 14 PASEF MS/MS scans were triggered per cycle (2.57 s) with a maximum of seven precursors per mobilogram. Precursor ions in an *m*/*z* range between 100 and 1700 with charge states ≥3+ and ≤8+ were selected for fragmentation. Active exclusion was enabled for 0.4 min (mass width 0.015 Th, 1/*k*_0_ width 0.015 V s/cm^2^). Protein identification and label free quantification^28^ was carried out using Peaks Studio X (Bioinformatics solutions Inc.). All proteomics in this manuscript were performed at the UC San Diego Biomolecular and Proteomics Mass Spectrometry Facility (BPMSF), and the facility is supported by the NIH shared instrumentation grant numbers S10 OD016234 (Synapt-HDX-MS) and S10 OD021724 (LUMOS Orbi-Trap).

Peptide peak area files were analyzed in R. Kinase motifs were determined using the Kinase Library^29^. Peak area files and analysis workspaces will be made available upon request. Raw data will be uploaded to the PRIDE server.

#### Western blotting and immunoprecipitation

Cell pellets for Western blot were collected by spinning cells at up to 1500 rcf in a centrifuge cooled to 4°C, and then the pellets were stored at −80°C prior to lysis. Lysis buffer was prepared by supplementing RIPA buffer with cOmplete protease inhibitor cocktail and PhosSTOP phosphatase inhibitor at the manufacturer’s recommended concentrations. For each 1 million cells in the cell pellets, 25 μL lysis buffer was added to cell pellets, gently mixed, and left on ice for 10 minutes. The lysate was then centrifuged at 18,000 rcf at 4°C for 10 minutes. The supernatant was removed and mixed with an appropriate volume of 4x Laemmli buffer and 2-mercaptoethanol (final concentration 2.5% v/v). The solution was mixed and then boiled for 10 minutes. After cooling, samples were loaded onto 4-15% gradient acrylamide gels and subjected to electrophoresis at 100V for approximately 1 hour. Proteins in the gel were transferred to a PVDF membrane in Towbin buffer at 100V for 90 minutes. Following transfer, membranes were blocked in a solution of 5% w/v BSA in TBST buffer at room temperature for 1 hour. The membranes were blotted with primary antibodies overnight at 4°C, rinsed 3 times in TBST at room temperature for 10 minutes and then blotted with an appropriate horseradish peroxidase (HRP) conjugated secondary antibody at room temperature for 1 hour. The membranes were rinsed in TBST for 10 minutes 3 times and then were imaged via chemiluminescence. Protein content was normalized to the loading control in each lane.

### QUANTIFICATION AND STATISTICAL ANALYSIS

#### Cell biology and in vivo experiments

Statistical significance of bar graphs and dot plots was assessed with Student’s t-test (2 conditions) or one way ANOVA (3 or more conditions). Kinetic experiments (T cell co-transfer experiments in Cl 13, tumor growth curves) were assessed by t-test or one way ANOVA at each timepoint as appropriate. Data are represented in figures as mean ± s.e.m. *Phosphoproteomics*: Peptide sequences and their peak areas were extracted from PEAKS software. Analysis of peak area data were carried out in RStudio as follows. Non-phospho-peptides were removed from the dataset to focus analysis on phosphorylation. Differential phosphorylation was calculated as the log2 fold change (log2FC) between two experimental conditions, e.g. anti-CD3 stim of *sgPrkcq* cells vs anti-CD3 stim of *sgPrkch* cells. These log2FC values for each peptide were then normalized by the log2FC of the unstimulated conditions for the same genotypes, e.g. unstimulated *sgPrkcq* vs *sgPrkch*, to account for proteomic drift during expansion of the distinct knockout genotypes. The normalized log2FC values for each phospho-peptide were then used as input for the Enrichment analysis feature of the Kinase Library software^29^ with default settings.

## Acknowledgements

We gratefully acknowledge all members of the Kaech lab for their generous help and suggestions and the Salk Flow Cytometry Core Facility for indispensable assistance and support. T.H.M. acknowledges the Damon Runyon Cancer Research Foundation for fellowship support.

## Declaration of Interests

T.H.M. and S.M.K. are coinventors on a provisional patent related to use of degradation-resistant PKC theta or perturbations of CK1G2 activity in adoptive cell transfer therapy. S.M.K. is a scientific advisory board member for Pfizer, EvolveImmune Therapeutics, Arvinas and Affini-T, and an Academic Editor at the *Journal of Experimental Medicine*. T.H.M., J.F., V.T., M.L., and S.M.K have conducted sponsored research in partnership with Arvinas.

