## Supplemental Materials for "Different signaling interpretations by PKC eta and theta control T cell function and exhaustion"

**Supplemental Information**

**Figure S1**

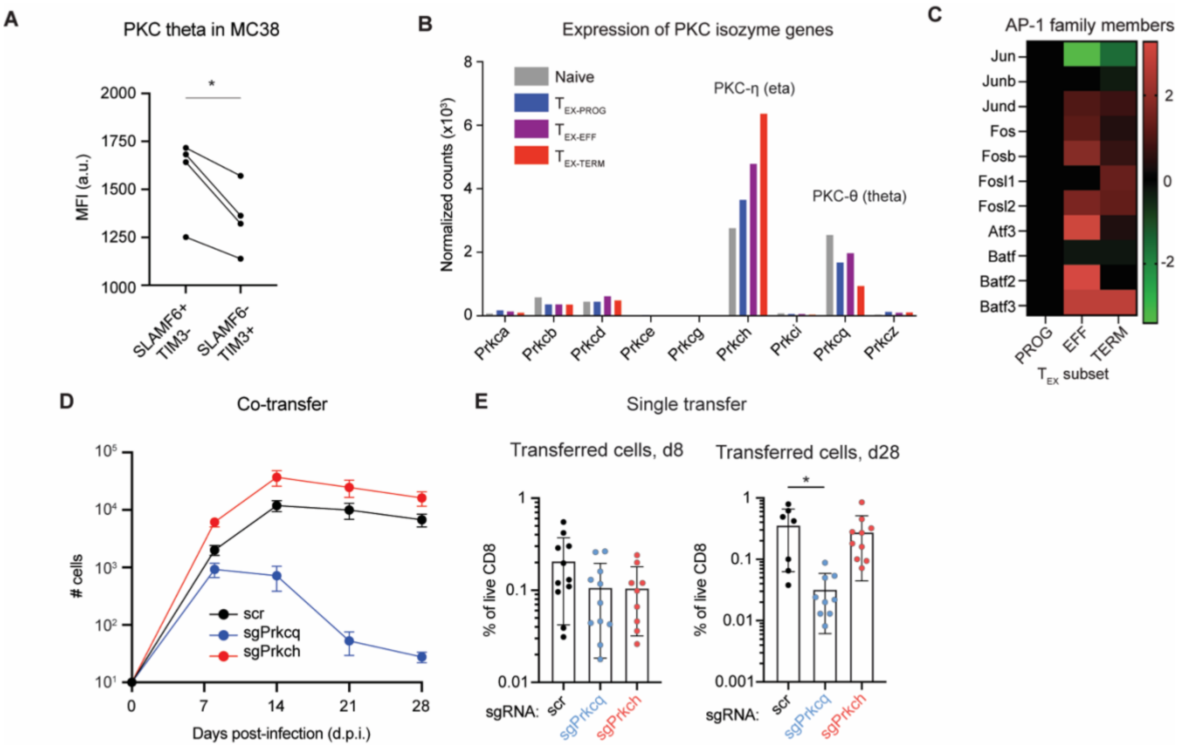


**Figure S1. Changes in the levels of PKC theta and eta mRNA and protein levels reflect different roles in $T_{EX}$ cell fate programming, related to Figure 1.**

(A) Anti-tumor CD8 T cells were isolated from MC38 tumors and stained for markers of $T_{EX}$ cell subset and for PKC theta.

(B-C) Expression of different PKC-encoding genes (B) or AP-1 factor genes (C) within sorted $T_{EX}$ subsets, reproduced from Hudson et al.[14]

(D) P14 T cells were electroporated (24 hours post-activation) with complexes of Cas9 bound to guide RNAs against *Prkcq*, *Prkch*, or a scrambled (scr) sequence control. After another 24 hours (48 hours post-activation), the electroporated T cells were mixed in approximately equal (1:1:1) proportions and adoptively transferred into mice that had been infected with LCMV Clone 13 48 hours prior. The mean number of T cells of each genotype present in the spleens is plotted by genotype over a 28-day timecourse.

(E) Similar experiments as in D were performed, but a single genotype of P14 cells was transferred into recipient mice immediately after Cas9 electroporation. The abundance of each genotype is indicated at 8- or 28-days post-infection as a fraction of the total live CD8 population in the spleen.

**Figure S2**

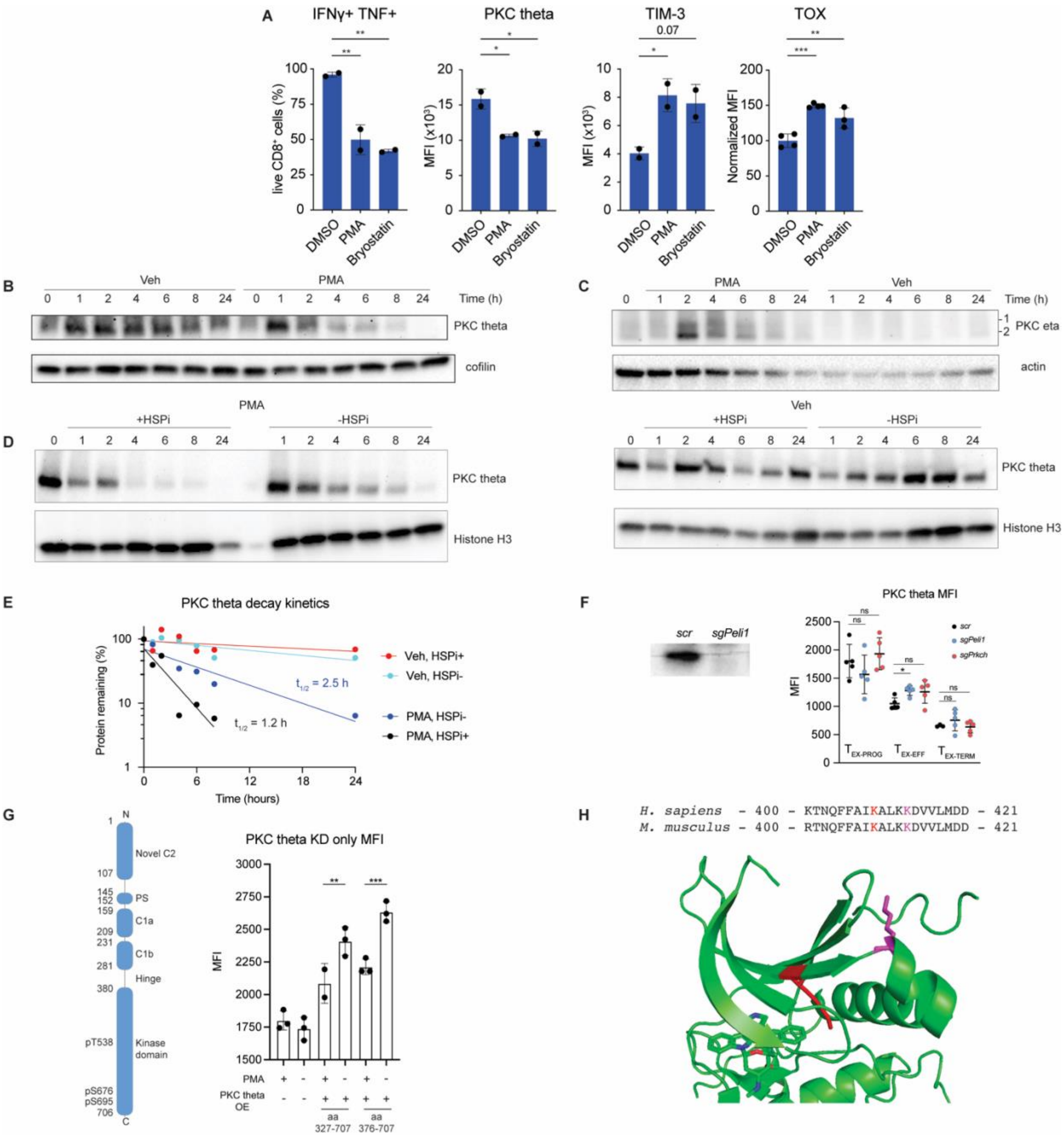


**Figure S2. Links between PKC theta agonism, degradation, and T cell exhaustion, related to Figure 2.**

(A) *In vitro* exhaustion phenotypes comparing PMA and bryostatin as PKC agonists.

(B-C) Western blot timecourse of PKC theta (B) and PKC eta (C) levels during PMA or vehicle treatment.

(D) Western blots of PKC theta levels in the presence of HSP70 inhibition.

(E) Degradation kinetics and half-lives ($t_{1/2}$) of PKC theta in presence of PMA or vehicle and in the presence of an HSP70 inhibitor (HSPi) or vehicle.

(F) Deletion of *Peli1*, encoding the E3 ligase for PKC theta, or PKC eta, and their effects on PKC theta protein levels by subset in P14 T cells responding to LCMV Clone 13 at 28 d.p.i. A Western blot shows validation of loss of the PELI1 protein.

(G) The kinase domain of PKC theta can be targeted for degradation during PMA stimulation. Two different variants of the catalytic C-terminus of PKC theta were overexpressed in $CD8^+$ T cells, either including the hinge region (aa 327-706) or lacking it (aa 376-706). T cells were treated with PMA or a vehicle control for 24 hours before being prepared for flow cytometry. The PKC theta domain architecture with key residues marked is reproduced from Figure 2.

(H) The crystal structure of the catalytic domain of human PKC theta is shown (PDB entry 1XJD). K409 is shown in red pointed towards the active site and K413 is shown in magenta.
